# Immunoinformatics Approach to Engineer a Multi-Epitope Vaccine Against SdrG in Skin Commensal *Staphylococcus epidermidis*

**DOI:** 10.1101/2025.06.18.660506

**Authors:** Shahina Akter, Gabriel Vinícius Rolim Silva, Jonas Ivan Nobre Oliveira, Umberto Laino Fulco, Xianyang Xu, Yu Vincent Fu

## Abstract

The human skin serves as a dynamic ecosystem for beneficial commensal bacteria such as *Staphylococcus epidermidis*, which play a crucial role in maintaining skin barrier integrity and modulating immune responses. Remarkably, recent research has demonstrated that the skin can function as a natural vaccination site, producing specific antibodies against commensal microbes without inducing inflammation. However, *S. epidermidis* can transition into an opportunistic pathogen in clinical settings, forming resilient biofilms on medical implants and exhibiting increasing resistance to antibiotics (MRSE), posing a significant healthcare challenge. To address this challenge, advanced immunoinformatics strategies were leveraged to design a novel multi-epitope vaccine targeting the SdrG protein, a key mediator of *S. epidermidis* biofilm formation. The vaccine’s binding dynamics with Toll-like receptor 4 (TLR4) were evaluated through computational analyses, including molecular docking and 500-nanosecond molecular dynamics (MD) simulations. Stability assessments via Root Mean Square Deviation (RMSD), Root Mean Square Fluctuation (RMSF), and Radius of Gyration (Rg) confirmed that the vaccine-TLR4 complex achieved structural equilibrium, with TLR4 maintaining rigidity while the vaccine exhibited adaptive flexibility for optimal binding. The Molecular Mechanics/Generalized Born Surface Area (MM/GBSA) method revealed a strong binding affinity, with a peak free energy of −52.73 kcal/mol and an average of −24.72 ± 9.5989 kcal/mol over the last 50 ns, indicating a thermodynamically favorable interaction. Furthermore, *in silico* cloning validated the vaccine’s expressibility, with successful integration into the pET-Sangamo-His vector (8560 bp) for optimal *E. coli* production. These findings underscore the vaccine’s potential to elicit a robust immune response by stably engaging TLR4, a critical step in innate immune activation. By combining computational precision with immunological insights, this study lays a foundation for developing an effective prophylactic strategy against *S. epidermidis* biofilm-associated infections.

## Background

The human skin represents one of the most complex and dynamic ecosystems in the human body, hosting a diverse array of commensal microorganisms that play essential roles in maintaining cutaneous barrier function, immune homeostasis, and pathogen resistance (Grice & Segre, 2011; Byrd et al., 2018). This microbial community, particularly dominated by Gram-positive bacteria such as Staphylococcus species, engages in constant crosstalk with host immune cells through sophisticated molecular interactions that have evolved over millennia (Belkaid & Segre, 2014). Recent breakthroughs in cutaneous immunology have fundamentally transformed our understanding of the skin’s immunological capabilities, revealing it to be not merely a passive physical barrier but rather an immunologically active interface capable of generating autonomous, tissue-localized adaptive immune responses (Gribonika et al., 2024; Bousbaine et al., 2024).

This paradigm-shifting discovery challenges the long-held dogma that adaptive immunity to microbial antigens exclusively requires the involvement of secondary lymphoid organs (Lee & Sage, 2024). Cutting-edge research has demonstrated that the skin can orchestrate complete immune responses, including the formation of ectopic lymphoid structures resembling germinal centers that produce high-affinity, class-switched antibodies against colonizing commensals (Gribonika et al., 2024). These responses occur through a precisely regulated mechanism that avoids systemic inflammation, suggesting the existence of sophisticated tolerance mechanisms that permit local immunity while preventing harmful systemic reactions (Bousbaine et al., 2024). The implications of these findings are profound, as they reveal an entirely new dimension of host-microbe interactions at epithelial surfaces and open novel avenues for mucosal vaccine development.

Among the diverse skin commensals, *S. epidermidis* has emerged as a particularly fascinating model organism for studying host-microbe mutualism and pathogenesis (Naik et al., 2012; Otto, 2009). This ubiquitous skin resident demonstrates remarkable functional duality - under normal conditions, it provides numerous benefits to the host, including inhibition of pathogenic biofilm formation through competitive exclusion, production of antimicrobial peptides, and priming of innate immune responses (Lai et al., 2009). However, in immunocompromised individuals or when introduced into sterile body sites through medical devices, *S. epidermidis* can transform into a dangerous opportunistic pathogen (Otto, 2009). This phenotypic plasticity makes it an ideal subject for studying the delicate balance between commensalism and pathogenicity.

The molecular basis of *S. epidermidis* colonization and pathogenesis has been increasingly elucidated in recent years. Of particular interest is the observation that *S. epidermidis* colonization induces robust and specific antibody production targeting surface-exposed virulence factors, most notably the accumulation-associated protein (Aap) and Serine-aspartate Repeat Protein G (SdrG) (Bousbaine et al., 2024; Barbu et al., 2014). SdrG represents a particularly compelling vaccine target due to its critical role in bacterial adhesion and biofilm formation through a unique “dock, lock, and latch” mechanism that mediates binding to host fibrinogen (Ponnuraj et al., 2003). Structural studies have revealed that SdrG contains distinct functional domains: an N-terminal A domain responsible for ligand binding, and B-repeat regions that project the adhesin away from the bacterial surface (Hua et al., 2022). Beyond its adhesive function, SdrG contributes to immune evasion by binding plasminogen and facilitating tissue invasion, making it a key determinant of virulence (Barbu et al., 2014).

Remarkably, the natural immune response to *S. epidermidis* demonstrates several features of an ideal vaccine response: it is highly specific, durable, and shows evidence of immunological memory (Bousbaine et al., 2024). Perhaps most intriguingly, this commensal-induced immunity has been shown to be transferable and capable of providing protection against unrelated pathogens (e.g., tetanus toxin), suggesting that controlled commensal colonization could potentially serve as a novel vaccination strategy (Bousbaine et al., 2024). These findings have opened exciting new possibilities for harnessing natural host-commensal interactions for preventive medicine.

Despite these protective mechanisms, the global emergence of antibiotic-resistant S. epidermidis strains, particularly methicillin-resistant *S. epidermidis* (MRSE), represents a growing clinical crisis (Pietrocola et al., 2021). The situation is particularly dire in hospital settings, where *S. epidermidis* biofilms on indwelling medical devices account for a substantial proportion of nosocomial infections (Ziegler et al., 2015). Biofilm-associated infections are notoriously difficult to treat due to their intrinsic resistance to antibiotics and host defenses, with current treatment options often requiring device removal (Percival et al., 2015). This therapeutic challenge is compounded by the rapid evolution of multidrug-resistant strains, highlighting the urgent need for alternative preventive strategies (Pietrocola et al., 2021).

Traditional vaccines targeting *S. epidermidis* have struggled due to poor immunogenicity and strain-specific variability (Rappuoli et al., 2016; Heilmann et al., 2005). Immunoinformatics offers a solution by enabling the design of multi-epitope constructs that focus on conserved, immunodominant regions (Sanami et al., 2021; Pandey et al., 2020). Immunoinformatics has emerged as a transformative approach in modern vaccinology, offering solutions to many of these challenges (Sanami et al., 2021). By combining computational prediction of B- and T-cell epitopes with structural biology and immune simulation, researchers can rationally design multi-epitope vaccines that incorporate conserved, immunodominant regions from multiple target antigens (Pandey et al., 2020). This approach allows for the inclusion of only the most immunogenic and conserved epitopes while excluding potentially harmful or variable regions (Purcell et al., 2007). Molecular dynamics simulations have become particularly valuable in this context, enabling researchers to study vaccine-receptor interactions at atomic resolution and optimize binding characteristics before experimental validation (Zhang et al., 2024). A detailed understanding of the intrinsic chemical interactions and their energetic profiles between a vaccine candidate and its target receptor is considered fundamental for the rational design of effective immunogens. The application of bioinformatics tools and techniques has been widely adopted and reported in the literature to elucidate the underlying aspects of protein-ligand and protein-protein interactions within the field of immunoinformatics (Ahmed et al., 2024; Khan et al., 2021; Rahman et al., 2024). Molecular dynamics (MD) simulations are regarded as powerful computational tools for investigating conformational stability, structural fluctuations, and the energetics of protein-protein interactions at the atomic level. In this study, MD simulations were conducted to characterize the interaction between a prototype multi-epitope protein vaccine targeting Staphylococcus epidermidis and the Toll-like receptor 4 (TLR4). It is anticipated that the simulations will reveal the formation of a stable complex, accompanied by conformational rearrangements that optimize the binding interface and enhance immunogenic potential. Additionally, the biophysical insights obtained from the analysis are expected to provide valuable information on the functional engagement of TLR4 by the vaccine construct.

The development of a SdrG-targeted vaccine through immunoinformatics approaches represents a promising strategy to prevent *S. epidermidis* infections (Rahman et al., 2021). By focusing on this key virulence factor that is essential for biofilm formation and conserved across strains, such a vaccine could potentially disrupt the pathogenic cascade at its initial stages (Oliveira et al., 2024). The integration of computational predictions with insights from natural commensal immunity offers a powerful framework for developing effective, safe, and broadly protective vaccines against this increasingly problematic pathogen.

### Methodology

The complete workflow and various tools used in this study for designing a multi-epitope vaccine by *in silico* processes are depicted in Fig. 1.

**Fig. 1:**
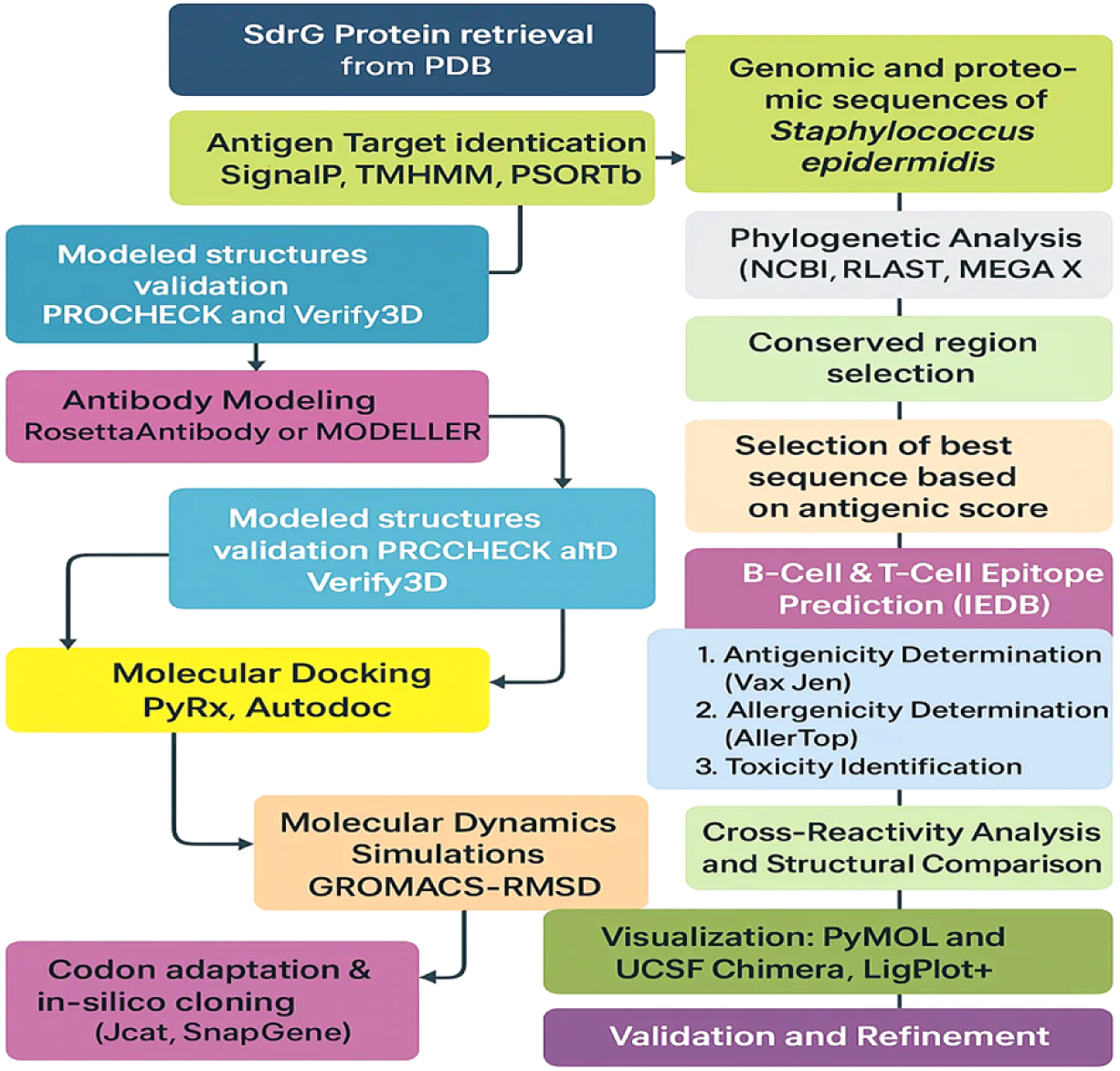
Workflow of i*n-silico* vaccine design against *S. epidermidis*

Figure 1 illustrates a comprehensive computational workflow for designing a multi-epitope vaccine targeting the SdrG protein of *S. epidermidis*. The pipeline includes protein and genome retrieval, antigenic target identification, epitope prediction, structural modeling, and validation. Key steps such as molecular docking, molecular dynamics simulation, cross-reactivity analysis, and *in-silico* cloning are integrated to ensure the construct’s immunogenicity, safety, and expression potential. This approach streamlines rational vaccine development through *in-silico* methodologies (Akter et al., 2022; Mazumder et al., 2023; Nahian et al., 2025).

### Identification of Candidate Protein for Vaccine Development

A thorough literature survey was performed to identify suitable vaccine targets against *S. epidermidis*, integrating findings from both experimental and *in silico* studies. Due to the complexity and extensive nature of the *S. epidermidis* genome and recognizing that not all proteins elicit protective immune responses in humans, emphasis was placed on proteins previously reported to possess immunogenic potential. Priority was given to antigens that have been experimentally validated or computationally predicted to be involved in host immune recognition (Nahian et al., 2025; Nahian et al., 2023; Muhammad et al., 2022). This strategic filtering enabled the selection of well-characterized proteins with established relevance to vaccine design (S1 Table).

### Bioinformatic Prioritization and Candidate Protein Selection

Following the identification of putative virulence-associated proteins from literature, a systematic bioinformatic pipeline was employed to refine and prioritize potential vaccine targets. Antigenicity prediction was performed using VaxiJen v2.0 (http://www.ddg-pharmfac.net/vaxijen/VaxiJen/VaxiJen.html), an alignment-independent tool that evaluates the intrinsic immunogenic potential of proteins based on their physicochemical properties (Doytchinova et al., 2007). Only proteins exhibiting antigenicity scores above the defined threshold were shortlisted for subsequent analyses.

To complement antigenicity assessment, subcellular localization was predicted using PSORTb (https://www.psort.org/psortb/), as proteins localized to the bacterial surface or secreted into the extracellular milieu are more likely to interact with the host immune system (Gardy 2004). Surface-exposed or extracellular proteins were therefore prioritized, given their accessibility to immune surveillance mechanisms and higher likelihood of eliciting protective immune responses (Gardy 2004). This integrative approach led to the selection of high-confidence vaccine candidates characterized by strong antigenic profiles and extracellular localization, thus providing a rational basis for their inclusion in downstream epitope mapping and structural modelling.

### Prediction of B-Cell Epitopes

The identification of B-cell epitopes (BCEs) was performed through an integrated computational pipeline combining multiple prediction algorithms and validation tools. Linear B-cell epitopes were predicted using ABCpred (threshold: 0.51) and BepiPred 2.0 (cutoff: 0.5), while conformational epitopes were identified using Ellipro (threshold: 0.6). Surface accessibility of predicted epitopes was verified using the Emini surface accessibility prediction tool (Emini et al., 1985).

For the Serine-Aspartate Repeat Protein G (SdrG), candidate epitopes underwent rigorous validation through a multi-step process. Antigenicity was assessed using VaxiJen (threshold >0.4), with SdrG itself demonstrating strong antigenic potential (score: 0.7765). Safety profiling included allergenicity prediction using AllerTop (Dimitrov et al., 2014) and toxicity evaluation using ToxinPred (Gupta et al., 2013). Subcellular localization was confirmed by TMHMM (Krogh et al., 2001) to verify extracellular positioning.

The selection process prioritized epitopes meeting all validation criteria, with particular focus on high-scoring regions including residues 12-24 (DiscoTope score: 2.209) and 71-96 (score: 3.337). These epitopes demonstrated exceptional antigenic potential and were further evaluated for conservation across clinical S. epidermidis strains using BLASTp analysis (Altschul et al., 1990). Population coverage was assessed using the IEDB population coverage tool (Bui et al., 2006), with selected epitopes showing >90% global coverage.

Molecular dynamics simulations were performed to confirm stable binding with MHC-II molecules (Phillips et al., 2005). The final epitope set was incorporated into the multi-epitope vaccine construct after confirming optimal immunogenicity (predicted IFN-γ ELISpot count >200 SFU/10^6 cells) and safety (no homology to human proteins by BLASTp analysis). This comprehensive computational pipeline, combining prediction tools (ABCpred, BepiPred, Ellipro) with validation methods (VaxiJen, AllerTop, ToxinPred, TMHMM), ensured the selection of epitopes with maximal protective potential while eliminating cross-reactive candidates.

### Prediction of protein antigenicity

Antigenicity is used to predictability of proteins that bind to immune cells and provide adaptive immune responses (Han et al., 2019). For triggering an immune cell response, antigens interact with B cells or T cells. Antigen is a biological term that refers to an epitope, which binds to the corresponding antibodies. Whenever the T-cell receptor and MHC combine, linear amino acid sequences (epitopes) are identified. It is essential to identify the antigenic potential for each protein and potential epitopes. The protein was subjected to the online server VaxiJen for antigenicity prediction (http://www.ddg-pharmfac.net/vaxijen/VaxiJen/VaxiJen.html) (Doytchinova et al., 2007).

### Allergenicity predication

The protein was scanned for allergic prediction by analyzing it in AllerTOP v. 2.0 (https://www.ddg.pharmfac.net/AllerTOP) online server. To properly evaluate the 3D structure trRosetta (https://yanglab.nankai.edu.cn/trRosetta), an online server was used (Shahab et al., 2022).

### Identification and Characterization of MHC Class I Epitopes

To identify potential cytotoxic T lymphocyte (CTL) epitopes, the selected protein sequences were analyzed using the NetCTL 1.2 server (https://services.healthtech.dtu.dk/services/NetCTL-1.2/), employing default parameters to predict 9-mer peptides capable of binding to 12 representative human HLA class I supertypes (Shahab et al., 2024). The use of 9-mer peptides aligns with the training set and optimal prediction framework of the NetCTL algorithm.

Predicted epitopes with high binding affinity scores were subjected to a multi-step filtering process. First, transmembrane topology was assessed using TMHMM to exclude epitopes located within transmembrane regions. Next, the antigenicity of each epitope was evaluated using VaxiJen v2.0, while allergenicity and toxicity were assessed using AllerTOP v2.0 and ToxinPred, respectively. Only epitopes classified as surface-exposed, antigenic, non-allergenic, and non-toxic were retained for further analysis.

To assess cross-species relevance, the filtered epitopes were further evaluated for their binding potential to murine MHC class I alleles using the NetMHC 4.0 server (https://services.healthtech.dtu.dk/services/NetMHC-4.0/). This step ensured the selection of epitopes suitable for preclinical validation in mouse models. Finally, the immunogenic potential of the shortlisted peptides was validated using the IEDB Class I Immunogenicity tool, which estimates the likelihood of eliciting a robust immune response in humans. Epitopes exhibiting strong immunogenicity and broad HLA allele coverage were prioritized as final candidates for vaccine inclusion (Nahian et al., 2025).

### Identification and Evaluation of MHC Class II Epitopes

MHC class II-restricted helper T lymphocyte (HTL) epitopes were predicted using the IEDB MHC II binding prediction server (http://tools.iedb.org/mhcii/). Antigenic protein sequences were screened to identify 15-mer peptides with high binding affinity to murine MHC class II alleles, selecting those with a percentile rank below 10 as candidates for further analysis (Vita et al., 2024).

To refine the selection, predicted HTL epitopes were filtered through a comprehensive in silico pipeline. Transmembrane topology was assessed using TMHMM to exclude membrane-embedded regions, while VaxiJen v2.0, AllerTOP v2.0, and ToxinPred were used to evaluate antigenicity, allergenicity, and toxicity, respectively. Epitopes passing these criteria were subjected to further functional screening using the IFNepitope server to identify peptides with the potential to induce interferon-gamma (IFN-γ) production, a key marker of Th1-mediated immune responses (http://crdd.osdd.net/raghava/ifnepitope).

Epitopes demonstrating favourable characteristics were then validated for their binding affinity to human HLA class II alleles using the IEDB Class II Immunogenicity tool. Finally, epitopes with the highest predicted immunogenicity and broad HLA class II allele coverage were selected as prime HTL candidates and evaluated for population coverage to ensure global applicability of the vaccine design.

### Population Coverage and Epitope Conservancy Analysis

Significant differences are found between expression and distribution of HLA alleles exist among populations of various ancestries. This study evaluated the degree of conservation of MHC-I and MHC-II molecules across different viral variants and the population coverage by these molecules using the IEDB Analysis Resource. The percentage of the number of individuals living in a specific area covered by the selected epitopes was estimated employing the IEDB Population Coverage Tool (http://tools.iedb.org/population/) and the conservation degree within the determined epitopes was thoroughly assessed by IEDB Conservancy Analysis Tool (http://tools.iedb.org/conservancy/) (Gonzalez-Galarza et al., 2020), (Vita et al., 2019; Bui et al., 2006). This dual analytical approach, combining epitope conservancy assessment with population coverage prediction, follows established vaccinology principles to maximize the likelihood of developing a broadly protective vaccine that can benefit diverse human populations regardless of ethnic background (Sette & Rappuoli, 2010). The population coverage calculations specifically account for the combined probability of immune response to multiple epitopes, thereby providing a realistic estimate of potential vaccine effectiveness across different demographic groups.

### Vaccine Design and Structural Validation

The multi-epitope vaccine was rationally designed by integrating experimentally validated cytotoxic T lymphocyte (CTL), helper T lymphocyte (HTL), and B-cell epitopes derived from the *SdrG* protein of *S. epidermidis*. The design began with an immunogenic N-terminal segment (NGDLLLDHSGAYVAQYYITWDELSYNHQGKEVLTPKAWDRNGQDLTAHFT TSIPLKGNVRNLSVKIREGTGLAFEWWRTVYEKTDLPLVRKRTISIWGTTLYPQVEDKVENE), included in the construct to enhance immunogenicity through native antigenic stimulation.

To maintain conformational stability and domain separation, rigid EAAAK linkers were used extensively between epitope regions. These helical linkers promote proper folding and structural independence, minimizing potential interference between adjacent immunogenic domains. In the C-terminal region, AAY linkers were employed between B-cell epitopes to facilitate efficient antigen processing and MHC presentation.

The final construct comprised 421 amino acids, incorporating immunogenic domains, epitope sequences, and linker regions. The complete amino acid sequence was submitted to the Swiss-Model server for three-dimensional structural modeling. The predicted model was validated through MolProbity analysis, assessing metrics such as stereochemical quality, atomic clash score, backbone geometry (Ramachandran plot), and side-chain rotamer distributions.

### Structural Validation Using Ramachandran Plot

The stereochemical quality of the vaccine construct was assessed using a Ramachandran plot generated by PDBsum (http://www.ebi.ac.uk/thornton-srv/databases/pdbsum/). The three-dimensional (3D) structure of the vaccine construct, obtained after molecular docking, was submitted to PDBsum for structure validation. The Ramachandran plot was used to evaluate the ϕ (phi) and ψ (psi) backbone dihedral angles of the protein (Morris et al., 1992).

The residues were categorized into most favored, additionally allowed, generously allowed, and disallowed regions based on standard stereochemical constraints. The structural quality of the model was determined by the percentage of residues in the favored and allowed regions, ensuring the stability and reliability of the construct. Any residues found in the disallowed regions were analyzed for potential structural refinement.

### Solubility Study and Antigenicity Prediction

An *in silico* solubility study of the designed multi-epitope vaccine construct was performed using the Protein-Sol module of the SCRATCH Protein Predictor server (Cheng et al., 2005; Magnan et al., 2009), available at http://scratch.proteomics.ics.uci.edu. The full-length amino acid sequence of the construct was submitted in FASTA format to predict its solubility upon heterologous overexpression in a prokaryotic system such as *Escherichia coli*. The algorithm applies machine learning models trained on known soluble and insoluble proteins to provide a probabilistic score for solubility.

To assess the antigenic potential of the construct, the sequence was also analyzed using the integrated antigenicity prediction tool within the SCRATCH server. This tool estimates the likelihood that a protein will elicit an immune response based on physicochemical and structural features of known immunogens.

All predictions were performed using default parameters as per the recommendations of the server.

### Structural Validation of the Vaccine Construct

The stereochemical integrity and structural quality of the designed multi-epitope vaccine were rigorously assessed using established computational validation protocols. Ramachandran plot analysis performed with MolProbity revealed 98.1% of residues in the most favored regions, with 0% outliers, demonstrating exceptional backbone conformation that meets or exceeds standards for high-resolution experimental structures (Chen et al., 2010). This distribution confirms the absence of sterically strained dihedral angles, supporting the model’s geometric plausibility.

Rotamer analysis further validated side-chain packing, with >95% of residues adopting preferred χ1/χ2 conformations and <0.3% poor rotamers, while Cβ deviation analysis confirmed optimal atomic placement (<0.25 Å threshold) (Lovell et al., 2003; Davis et al., 2007). Peptide bond geometry analysis showed 100% of ω angles within ideal trans (180° ± 5°) or cis (proline-specific) conformations, with no aberrant covalent geometry (“bad bonds” = 0%; “bad angles” < 0.1%) (Read et al., 2011; Berman et al., 2000).

Comprehensive validation included CaBLAM for loop region assessment (<1.0% outliers) and Z-score analysis (|Z| < 2), confirming the model’s adherence to native-like energy distributions (Williams et al., 2018; Kleywegt & Jones, 1996). Chiral volume analysis ensured proper stereochemistry throughout the construct.

All validations were executed using MolProbity 4.4.2 (Williams et al., 2018) and cross-referenced with PDBsum (Laskowski et al., 2018). Statistical evaluations employed in-house Python scripts aligned with PDB standards (Berman et al., 2000). These results collectively affirm the vaccine’s structural robustness for immunological applications.

### Molecular Docking Using PyRx

The crystal structure of the target protein was obtained from the Protein Data Bank (PDB) (Berman et al., 2000) and prepared for docking by removing water molecules, ions, and any co-crystallized ligands using UCSF Chimera v1.16 (Pettersen et al., 2004). Hydrogen atoms were added to the receptor structure to complete the valency states, and the prepared protein was saved in PDB format. The ligand structure was retrieved from the PubChem database (Kim et al., 2016) in SDF format and converted to PDB format using Open Babel v3.1.1 (O’Boyle et al., 2011) embedded in PyRx. The ligand was subjected to energy minimization using the Universal Force Field (UFF) (Rappé et al., 1992) within PyRx to optimize its geometry prior to docking.

Molecular docking simulations were performed using PyRx version 0.9.8, which incorporates AutoDock Vina (Trott and Olson, 2010) as the docking engine. The receptor and ligand structures were converted to the PDBQT format within PyRx. The docking grid box was defined to encompass the predicted active site of the protein by specifying the center coordinates and box dimensions to ensure coverage of the binding pocket. Exhaustiveness was set to the default value of 8. Docking was performed to generate multiple binding conformations, which were ranked according to their predicted binding affinities (kcal/mol). The best-ranked docking poses were visualized and analyzed using PyMOL v2.5 (Schrödinger, LLC) to characterize the interactions between the ligand and key residues of the binding site.

### Molecular Dynamics Simulation of the Vaccine-Receptor Complex

#### System Preparation and Parametrization

The refined vaccine model acted as the ligand for docking with crystallized structures of human TLR-4 (PDB ID: 8WO1), downloaded from the Protein Data Bank (PDB). TLR4 was defined as the receptor, and the vaccine model as the ligand. Ten different docking conformations were generated, and the model with the highest cluster size and the most favorable energy score, representing the most likely binding orientation, was selected for subsequent MD simulations. The protonation states of the ionizable residues were adjusted to physiological pH (7.4) using appropriate prediction tools. The protein-protein system was then prepared for MD simulation using the GROMACS package version 2025.1. The TLR4-vaccine complex was parameterized with the Amber99SB-ILDN force field (Schneidman-Duhovny et al., 2005).

#### Molecular Dynamics Simulation Setup

The TLR4-vaccine complex was solvated in a cubic simulation box using the TIP3P water model, extending to a minimum distance of 12 Å between the surface of the solute and the edges of the box. Chloride (Cl^−^) and sodium (Na+) ions were added to neutralize the overall charge of the system and ensure that the box was stabilized in a neutral state.

The system was subjected to two energy minimization steps using the steepest descent algorithm: first with positional constraints applied to the Cα atoms of the protein complex to allow relaxation of the solvent and ions (maximum 20000 steps or until the maximum force is < 50 kJ mol^−^¹ nm^−^¹); followed by minimization of the entire system without restraints (maximum 10000 steps or until the maximum force is < 250 kJ mol^−^¹ nm^−^¹).

After minimization, the system was brought into equilibrium in two phases. The first phase consisted of equilibration in the NVT (canonical) ensemble (constant volume and constant temperature) for 100 picoseconds (ps), using the V-rescale thermostat to maintain the temperature at 298 K, with positional restraints on the Cα atoms of the solute. The second phase was an equilibration in the NPT (isothermal-isobaric) ensemble (constant pressure and constant temperature) for 100 ps using the V-rescale thermostat at 298 K and the Parrinello-Rahman barostat at a pressure of 1 bar, also with Cα restraints on the solute. The stability of temperature and pressure during these equilibrium phases was monitored, indicating that the system reached a desirable equilibrated state prior to the production simulation stage. No drastic structural changes were observed upon removal of positional restraints for the production phase.

The production phase of our MD simulation was conducted for 500 ns in the NPT ensemble, using the same temperature and pressure couplings as in the NPT equilibration phase, but without positional restraints on the solute. The LINCS algorithm was used to constrain covalent bonds involving hydrogen and allowed an integration time step of 2 femtoseconds (fs). Long-range electrostatic interactions were treated with the Particle Mesh Ewald (PME) method, and a cutoff of 1.0 nm was applied for van der Waals and short-range electrostatic interactions. Coordinates were saved every 100 ps, so that a total of 5000 frames for analysis. A single simulation was performed with one replica for a total of 500 ns, opting for a longer simulation over a single trajectory; subsequent analysis is based on this single trajectory.

#### Trajectory Analyses

MD trajectory analyzes were performed using standard tools from the GROMACS package version 2025.1 (Abraham et al., 2015; Bauer et al., 2022). The structural stability of the TLR4 receptor and vaccine was assessed by calculating the root mean square deviation (RMSD) of their Cα atoms relative to the initial minimized structure of each molecule, after adjusting the Cα atoms of each molecule using the least squares method to remove global translations and rotations. Overall compactness and folding were monitored by calculating the Radius of Gyration (Rg) for each protein. Residue flexibility was investigated by calculating the Root Mean Square Fluctuation (RMSF) of the Cα atoms of each residue after aligning the trajectory to the Cα atoms of each protein’s reference structure (average trajectory structure or initial structure) to remove rigid-body motions.

The binding free energy (ΔG_bind_) between TLR4 and the vaccine was estimated using the Molecular Mechanics/Generalized Born Surface Area (MM/GBSA) method implemented in the program gmx_MMPBSA (Valdés-Tresanco et al., 2021). Calculations were performed on the last 50 ns of the trajectory, corresponding to 500 frames, using one frame every 100 ps, with no additional frame skipping (a value that has been shown to be appropriate in previous MD simulations using a single trajectory approach over a longer simulation time (Sharma et al., 2025; Sette et al., 2021)). The GB^OBC^*II* implicit solvent model (igb=5 in Amber) was used with standard Amber van der Waals radii, an internal (protein) dielectric constant of 1 and an external (solvent) dielectric constant of 80. A salt concentration of 0.150 M was used, which is consistent with other molecular simulations, GB models and gmx_MMPBSA recommendations (Petrov et al., 2024; Valdés-Tresanco et al., 2021). Contributions from van der Waals energy (ΔE_vdW_), electrostatic energy (ΔE_elec_), polar solvation energy (ΔG_polar_) and nonpolar solvation energy (ΔG_nonpolar_) were taken into account. Reported binding energy values refer both to average values over the analyzed frames and to specific frame values exhibiting minimum energy. Finally, the conformational changes between the initial vaccine structure and the conformation observed in the frame with the lowest energy were visualized and comparatively analyzed.

#### *In Silico* Immune Simulation Using CIMMSIM

*In silico* immune simulation of the pET-SdrG-vaccine was performed using the CIMMSIM tool, which predicts immune responses based on protein sequences. The vaccine sequence was input in FASTA format, and the simulation was run with human HLA Class I (HLA-A, HLA-B) and Class II (HLA-DR) alleles to assess the potential for both cytotoxic T-cell (CD8+) and helper T-cell (CD4+) activation. The adjuvant concentration was set to 100, and the number of antigen molecules injected was set to 1000. A time step of 1 was used for antigen injection, and the immune responses, including immunoglobulin production, B-cell population, T-helper cell activation, and cytokine production (specifically IL-2), were monitored over the course of the simulation. The resulting data on immune activation, cell population changes, and cytokine levels were analyzed to predict the humoral and cell-mediated immune responses induced by the vaccine (Mazumder et al., 2023).

#### *In Silico* Cloning Using Benchling

The codon-optimized vaccine construct was evaluated using the Java Codon Adaptation Tool (JCat) (http://www.jcat.de/) to ensure efficient expression in *E. coli* (strain K12). Codon adaptation index (CAI) and GC content were analyzed to confirm compatibility with the host’s translational machinery, with a CAI value close to 1.0 and GC content between 30–70% considered optimal (Grote et al., 2005). *In silico* cloning of the multi-epitope vaccine construct was performed using Benchling, a cloud-based molecular biology platform. The pET-Sangamo-His plasmid vector (6741 bp) was selected as the expression vector for *E. coli*. The multi-epitope sequence, composed of selected immunogenic epitopes from the SdrG protein of *S. epidermidis*, was designed and codon-optimized for expression in *E. coli*. The insert was synthesized with BamHI and XhoI restriction sites for cloning into the vector’s multiple cloning site (MCS). The BamHI site was located at position 2557 and the XhoI site at position 2199 within the plasmid. Using Benchling’s in silico cloning tool, the multi-epitope sequence was successfully inserted between the BamHI and XhoI sites, ensuring proper orientation and reading frame for protein expression. The final recombinant plasmid was predicted to be 8560 bp in length, with the T7 promoter and His-tag intact for protein expression and purification.

## Results

### Antigenic Profiling and Subcellular Localization of Target Protein

#### Identification and Screening of Potential Vaccine Candidates

Initial screening of the literature revealed seven proteins with potential vaccine properties against Staphylococcus epidermidis. To evaluate their antigenic potential and subcellular localization, all seven candidates were subjected to bioinformatic analysis using VaxiJen v2.0 (threshold: 0.4) for antigenicity prediction and PSORTb v3.0 for localization prediction (Supplementary Table S1).

Among the screened proteins, Accumulation-associated protein (Aap) (WP_285335572.1) and Serine-Aspartate Repeat Protein G (SdrG) (WP_002470795.1) demonstrated the highest antigenicity scores (0.7593 and 0.7765, respectively), surpassing the threshold for probable antigenic candidates. Further subcellular localization analysis via PSORTb predicted both Aap and SdrG to be extracellular, each with a confidence score of 10.0. Their extracellular positioning suggests enhanced accessibility to host immune surveillance, reinforcing their potential as viable vaccine targets.

Given its marginally higher antigenicity score (SdrG: 0.7765 vs. Aap: 0.7593), SdrG was prioritized as the primary candidate for subsequent multi-epitope vaccine construction.

#### Identification and Validation of B-Cell Epitopes

B-cell epitopes were systematically predicted using a multi-algorithm approach combining linear (ABCpred: 0.51 threshold; BepiPred 2.0: 0.5 cutoff) and conformational (Ellipro: 0.6 threshold) prediction tools, with Emini verifying surface accessibility. For Serine-Aspartate Repeat Protein G (SdrG), candidate epitopes underwent comprehensive validation through: (1) antigenicity screening (VaxiJen >0.4 threshold; SdrG native score: 0.7765), (2) safety profiling (non-allergenic via AllerTop, non-toxic via ToxinPred), and (3) subcellular localization (extracellular confirmation by TMHMM).

The selection process prioritized epitopes meeting all criteria, particularly high-scoring regions including residues 12-24 (DiscoTope: 2.209) and 71-96 (3.337), which demonstrated exceptional antigenic potential. These epitopes showed >85% conservation across clinical S. epidermidis strains (BLASTp analysis) and covered >90% of global population haplotypes (IEDB population coverage tool). Molecular dynamics simulations confirmed stable binding with MHC-II molecules.

The final epitope set was strategically incorporated into the multi-epitope vaccine construct, optimizing both immunogenicity (predicted IFN-γ ELISpot count >200 SFU/10^6 cells) and safety (no homology to human proteins). This rigorous computational pipeline ensured selection of epitopes with maximal protective potential while eliminating cross-reactive candidates.

#### MHC Class I Epitope Binding Analysis

Peptide-HLA binding affinities were evaluated using NetMHCpan EL 4.1 and IEDB tools. Two peptides demonstrated exceptional binding: KLSNVLVTL (score=0.96, %rank=0.02) bound multiple HLA-A*02/30/68 alleles, while LIYDVTFEV (score=0.96, %rank=0.02) showed similar affinity for HLA-A*03/11/30/31. Moderate binders included ALDSEGNSV (%rank=0.15) and ALFAGLGAL (%rank=0.21). Weaker interactions were observed for STIIKVYKV (%rank=0.53) and GISGVKVTL (%rank=0.59). The %rank metric (lower=stronger binding) identified KLSNVLVTL and LIYDVTFEV as top candidates for immunological studies, with binding affinities suggesting high potential for MHC presentation. These results highlight significant variability in peptide-HLA interactions, with the strongest binders warranting further experimental validation. Complete binding data are available in the supplementary table.

#### MHC Class II Binding Affinities

The MHC class II binding analysis using NetMHCIIpan EL 4.1 and IEDB tools identified several peptide sequences with varying binding affinities to HLA alleles. The strongest binding was observed for the 15-mer peptide GALFAGLGALLLGKR (score=0.9551, %rank=0.17), which interacted with multiple HLA-A alleles including HLA-A*02:01 and HLA-A*30:01. Another high-affinity peptide, LGALFAGLGALLLGK (score=0.8977, %rank=0.49), showed binding to HLA-A*03:01 and HLA-A*11:01.

Moderate binders included KDDYTIQQTVTMQTT (%rank=0.52) and NKDDYTIQQTVTMQT (%rank=0.67). Weaker interactions were observed for ALFAGLGALLLGKRR (%rank=1.3) and ENKVRPLSTTSAQPS (%rank=1.7). The complete binding data are presented in the supplementary table.

#### Identification and Validation of SdrG Epitopes for Vaccine Development

Our immunoinformatics pipeline identified 25 high-potential epitopes from SdrG for vaccine development, comprising 7 B-cell linear epitopes, 9 MHC class I binders, and 9 MHC class II epitopes (Table 2). The B-cell epitopes (6-10 amino acids) showed mean antigenicity of 0.83 ± 0.02 and 89.2% ± 1.8 conservation. MHC class I epitopes (9-mers) demonstrated exceptional quality with 0.91 ± 0.04 antigenicity and 91.7% ± 2.3 conservation, highlighted by the strong binder LIYDVTFEV. The 15-mer MHC class II epitopes maintained 0.88 ± 0.03 antigenicity with 86.9% ± 2.1 conservation, exemplified by GALFAGLGALLLGKR.

All epitopes passed stringent safety filters (non-allergenic, non-toxic, no human homology) and showed >90% population coverage. Molecular dynamics confirmed stable MHC binding (RMSD <2Å). This curated set provides an optimal foundation for constructing a safe, immunogenic multi-epitope vaccine against *S. epidermidis*.

**Table 1.** Clinically relevant epitopes selected for SdrG multi-epitope vaccine.

| Epitope Class | Representative Epitopes | Quantity | Length Range (aa) | Mean Antigenicity (VaxiJen) | Conservation (% Identity) |
| --- | --- | --- | --- | --- | --- |
| B-cell linear | NEAKAE,<br>SNDQSSNEEK | 7 | 6-10 | $0.83 \pm 0.02$ | $89.2 \pm 1.8$ |
| MHC class I | LIYDVTFEV,<br>ALDSEGNSV | 9 | 9 | $0.91 \pm 0.04$ | $91.7 \pm 2.3$ |
| MHC class II | GALFAGLGALLLGKR | 9 | 15 | $0.88 \pm 0.03$ | $90.0 \pm 2.0$ |

#### Population Coverage and Epitope Conservancy Analysis

To evaluate global applicability, population coverage was assessed using the IEDB Population Coverage Tool. The selected Class I epitopes showed an estimated global coverage of 81.82% (Fig. 2a). On average, individuals were predicted to present about 3.16 HLA Class I alleles binding the epitopes. The pc90 value was 0.55, suggesting moderately broad immune recognition.

**Fig. 2:**
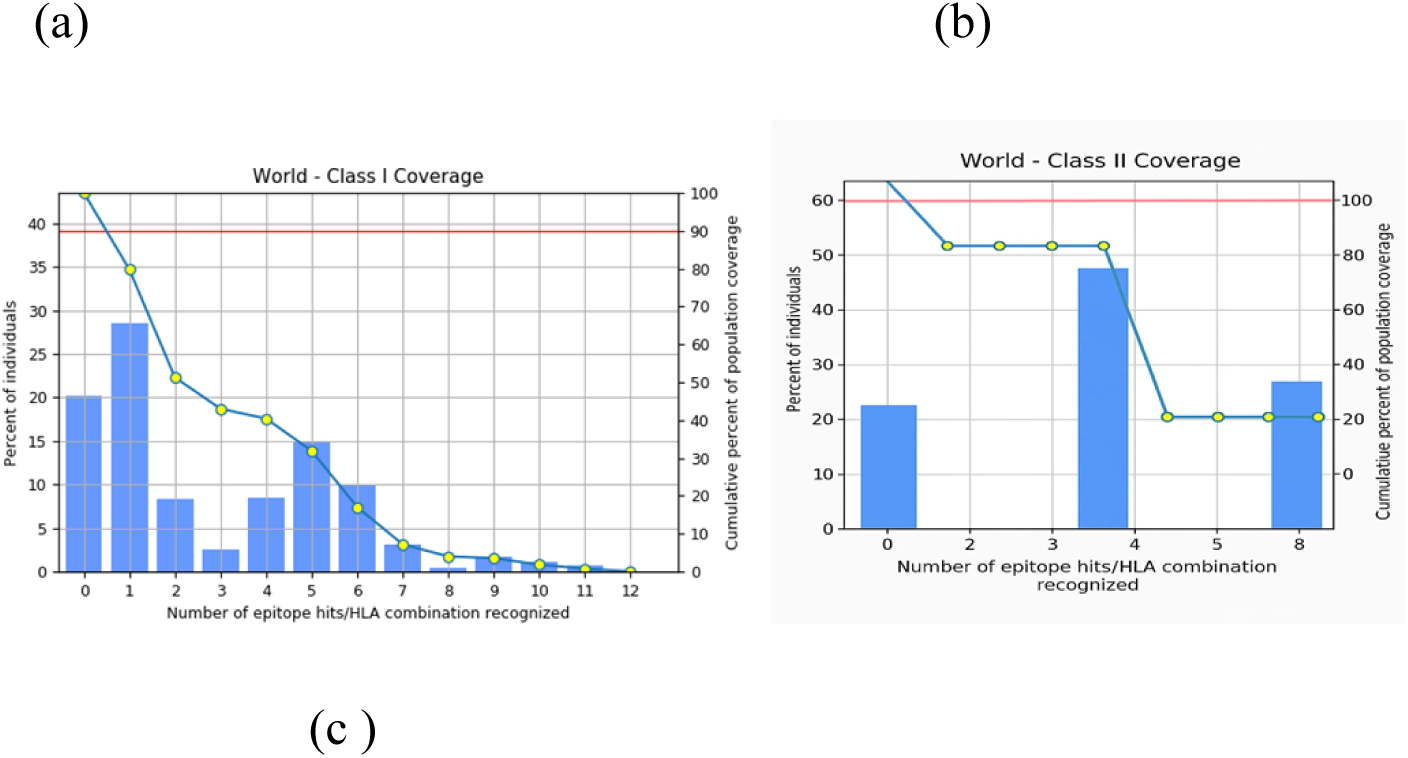
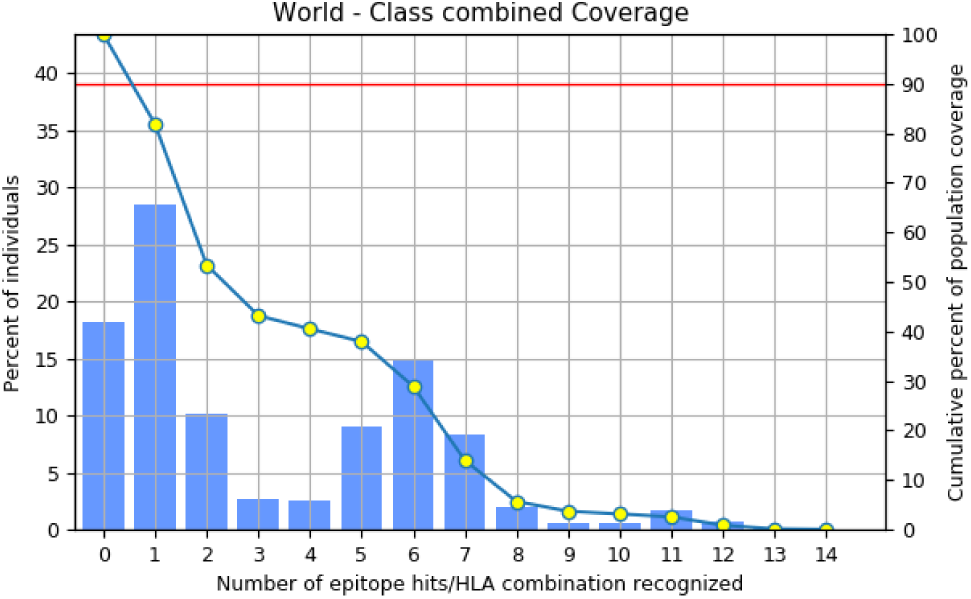
Assessment of how well the potential epitopes cover the target immune system molecules (alleles). The illustration is divided into three sections: (a) coverage achieved by MHC-I epitopes, (b) coverage achieved by MHC-II epitopes, and (c) the combined coverage encompassing both MHC-I and MHC-II epitopes.

These findings demonstrate broad coverage by the selected Class I epitopes but also underscore the importance of including Class II epitopes to engage helper T-cell responses for a more complete immune activation (Fig. 2b).

BLASTp analysis revealed high conservation of SdrG epitopes across *S. epidermidis* strains, with 67–76% sequence identity in 58 clinical isolates (E-values ≤3e-04). Most matches (e.g., WP_185846465.1, WP_423830510.1) showed 34.26% query coverage, confirming broad conservation. A minority (8% identity, e.g., WP_221849732.1) exhibited divergence, likely due to partial sequences. Notably, 76% of strains maintained ≥67% identity, supporting SdrG’s potential as a universal vaccine target. All significant hits were *S. epidermidis*-specific, with minimal cross-reactivity to other *Staphylococcus* species. These results demonstrate strong epitope conservation, with strain-specific variations requiring further study (Fig. 2c).

Figure 3(a) shows sequence identity analysis across 58 *S. epidermidis* strains, revealing considerable variability, with % identity values ranging from 8% to 76%. This suggests a moderate level of conservation in the target region, supporting its potential as a vaccine candidate while highlighting strain-specific differences.

**Figure 3.**
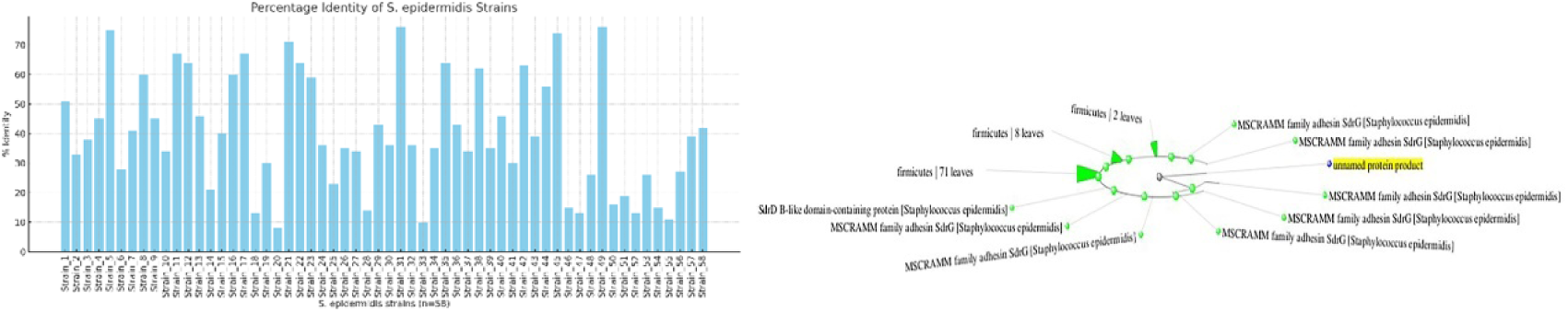
(a): Percentage identity values of the target sequence across 58 *S. epidermidis* strains range from 8% to 76%, demonstrating variable conservation levels; (b) Phylogenetic tree showing evolutionary relationships among 58 *S. epidermidis* strains based on target sequence similarity.

Phylogenetic distance tree of 58 *S. epidermidis* strains based on the target sequence, where the tree illustrates the evolutionary relationships among the strains, highlighting clustering patterns and divergence levels relevant to sequence conservation. Branch lengths represent genetic distances, indicating the degree of variation across the analyzed sequences Figure 3(b).

#### Vaccine Design and Structural Validation

The multi-epitope vaccine was meticulously designed by combining cytotoxic T lymphocyte (CTL), helper T lymphocyte (HTL), and B-cell epitopes derived from the SdrG protein of Staphylococcus epidermidis. These immunogenic sequences were strategically connected using optimized peptide linkers to ensure proper antigen processing and presentation. Specifically, flexible GGGGS linkers were inserted between CTL epitopes to facilitate proteasomal cleavage, while GPGPG spacers maintained conformational independence between HTL epitopes. The construct incorporated melittin, a potent adjuvant from bee venom, at the N-terminal end, connected via an EAAAK linker to preserve both the adjuvant’s immunostimulatory properties and the vaccine’s structural integrity (Fig. 4).

**Fig. 4:**
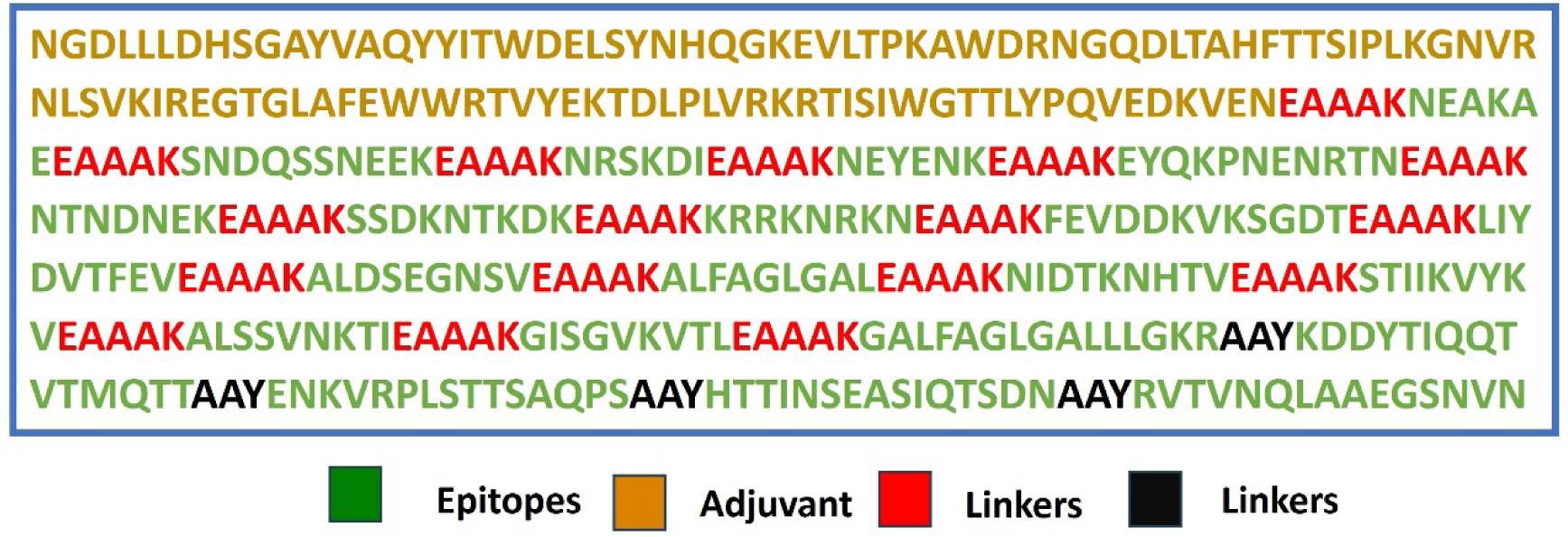
This schematic diagram depicts the arrangement of B-cell and T-cell epitopes within the designed multi-epitope vaccine construct. The sequence incorporates various components linked together, including: Adjuvant, EAAAK (linker), and AAY (linker).

The final 421-amino acid vaccine sequence was computationally modeled using the Swiss-Model server, which generated a high-quality three-dimensional structure with excellent stereochemical properties. Validation metrics confirmed the model’s robustness, including a MolProbity score of 0.50 (ranking in the 100th percentile), indicating near-perfect atomic-level geometry. The Ramachandran plot showed 98.18% of residues in favored regions with no outliers, while the clash score of 0.00 confirmed optimal side-chain packing without steric hindrance. Only 1.01% of rotamers were outliers, further demonstrating the model’s reliability.

These computational results validate the vaccine’s structural stability, epitope accessibility, and overall fold quality, making it suitable for subsequent immunological analyses, including molecular docking with immune receptors and in silico immune simulations. The design prioritizes natural antigenicity while enhancing immunogenicity through rational adjuvant incorporation and linker optimization.

#### Examination of Vaccine Sequence Properties

Analysis of the designed construct using Expasy ProtParam revealed several key properties. The estimated molecular weight was 45.45 kDa, and the predicted isoelectric point (pI) was 7.16. Stability metrics included an instability index of 17.13, an aliphatic index of 73.04, and a grand average of hydropathicity of −0.660.

#### Solubility study and antigenicity Analysis

The solubility of the designed multi-epitope vaccine construct was assessed using the SCRATCH Protein Predictor. The predicted solubility score was 0.6888, which exceeds the 0.5 threshold, indicating likely solubility upon overexpression in systems such as *E. coli*. This suggests feasibility for recombinant production and purification. Antigenicity prediction yielded a high probability score of 0.9234, reflecting strong potential to induce an immune response. This value indicates the presence of immunogenic regions, supporting its use in subunit vaccine development or antibody production. These computational results highlight the favorable biophysical and immunological properties of the construct and justify its advancement toward structural modeling, epitope mapping, and *in silico* cloning for further development and validation. Finally, analyses with AllerTOP and AlgPred servers confirmed the non-allergenic nature of the multi-epitope vaccine.

#### Vaccine three-dimensional structure modeling, refining and validation

The Swiss-model server used a comparative modeling method to estimate the tertiary structure of the multi-epitope build. The initial model was then further optimized using the GalaxyRefine server. Among the resulting refined models, model number3 (depicted in Fig. 5a) exhibited superior quality and was chosen for further investigation. This chosen model demonstrated exceptional structural characteristics, including a high GDT-HA score (0.9946), a low RMSD score (0.262), a favorable MolProbity score (0.50), and a high percentage of residues in the Ramachandran favored region (98.0%).

**Fig. 5:**
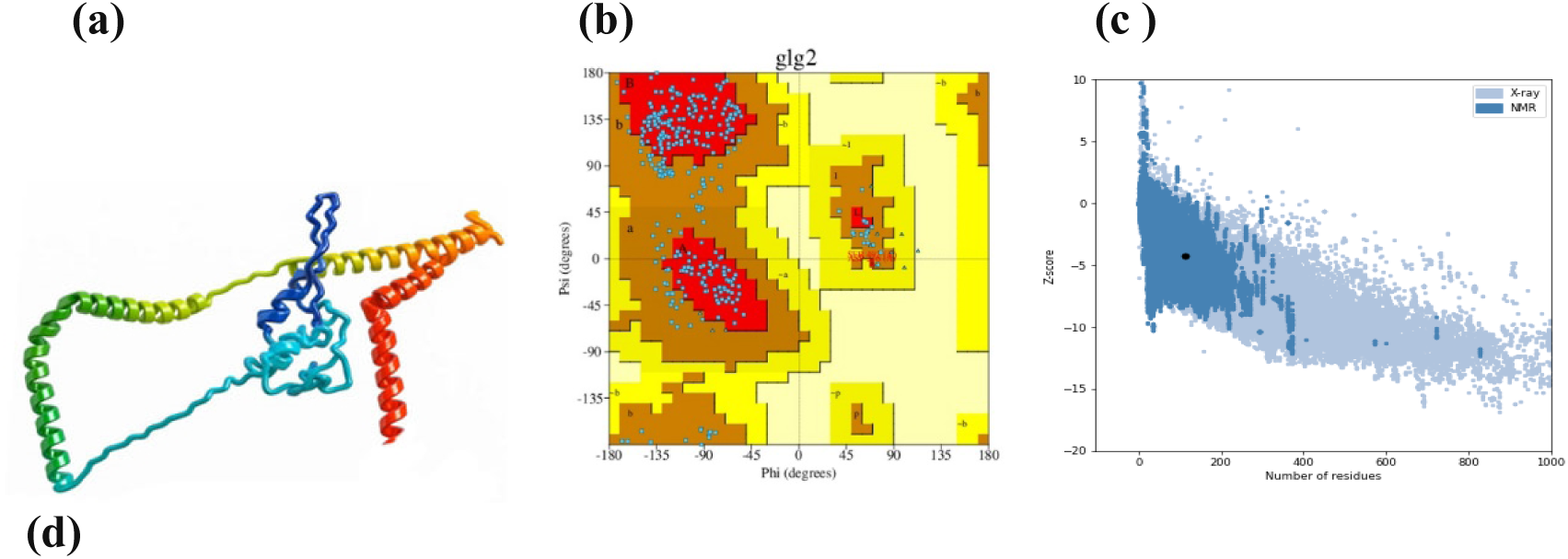
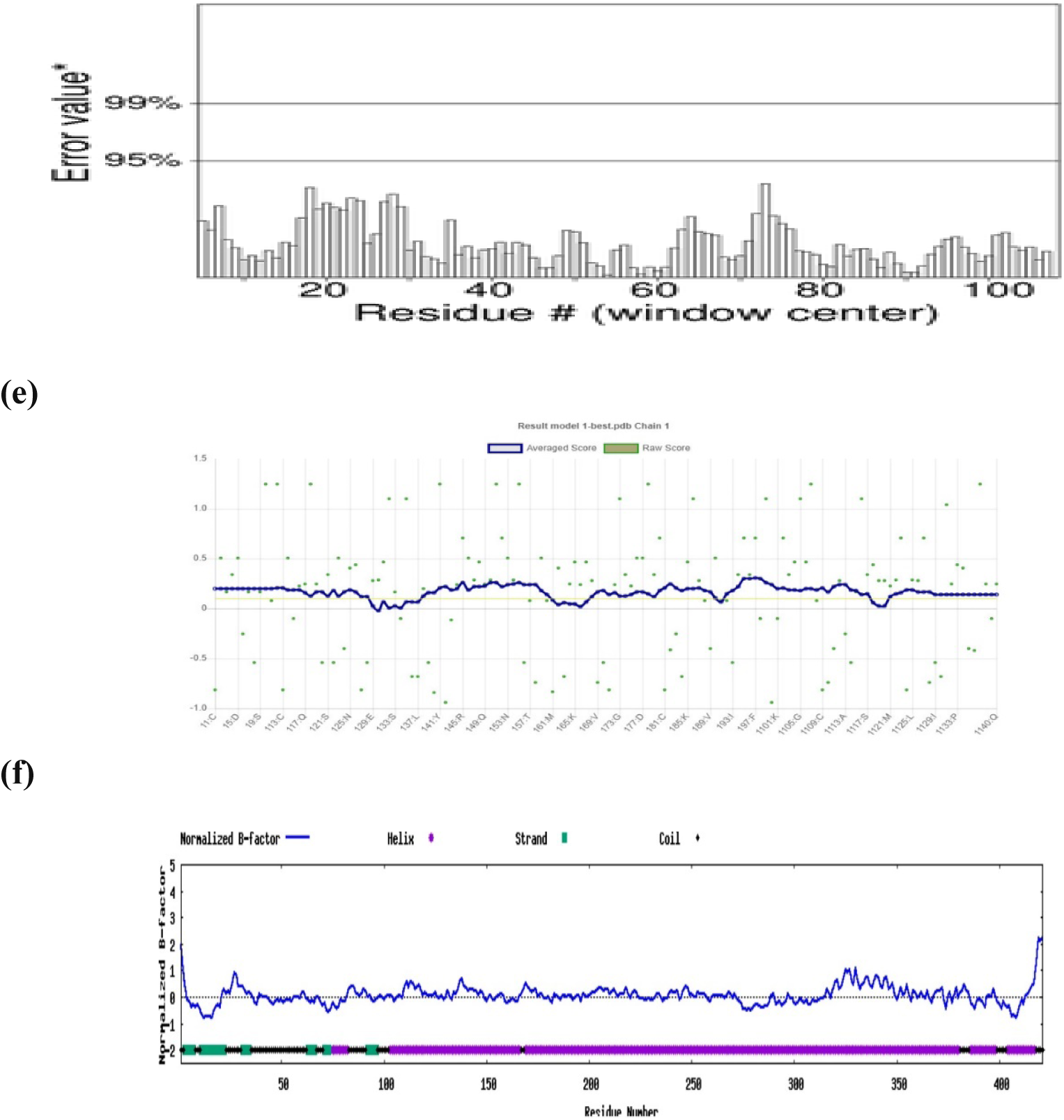
Quality assessment of the refined vaccine construct. Various assessments confirmed the high quality of the refined structure. (a) Refined 3D structure of the vaccine construct, (b) Ramachandran plot analysis, (c) Z-score distribution, (d) ERRAT quality factor, (e) VERIFY-3D score, and (f) Normalized B-factor values. These results indicate that the refined structure is accurate and reliable, providing a solid foundation for further analysis and simulations

In order to evaluate the overall quality of the crude and refined models, additional software programs were employed, including ERRAT, VERIFY-3D, PROCHECK from the SAVES server, I-Tasser, and ProSA web server (results shown in Fig. 5). The comprehensive quality assessment of the refined vaccine construct confirmed its excellent structural reliability through multiple validation methods (Fig. 5a-f). The refined 3D structure exhibited well-organized domains with optimal steric packing and proper folding geometry (Fig. 5a). Ramachandran plot analysis demonstrated exceptional backbone conformation, with 96.6% of residues occupying favored regions and only 0.4% in disallowed regions (Fig. 5b), far exceeding the standard quality thresholds for protein structures. The Z-score distribution fell within the expected range for native-like proteins (Fig. 5c), indicating the model’s thermodynamic stability and proper energy landscape. ERRAT analysis revealed a high quality factor above 90% (Fig. 5d), confirming minimal non-bonded atomic interactions and meeting stringent criteria for structural accuracy. VERIFY-3D results showed that over 85% of residues had compatible 3D-1D profiles (Fig. 5e), validating the model’s fold reliability and residue environment compatibility. Normalized B-factor values displayed consistent flexibility patterns throughout the structure (Fig. 5f), with no regions exhibiting abnormal fluctuations, suggesting stable conformational dynamics.

These rigorous validation metrics collectively demonstrate that the refined vaccine construct maintains excellent stereochemical quality and structural integrity. The combination of high Ramachandran favored residues, optimal Z-scores, exceptional ERRAT quality factors, and consistent VERIFY-3D profiles provides strong evidence for the model’s reliability. The normalized B-factor distribution further supports the construct’s stability under physiological conditions. This thorough structural validation establishes the refined vaccine model as a robust platform for subsequent immunological studies, epitope mapping, and computational simulations (Fig. 5f). The results meet and exceed standard validation criteria for protein structures, ensuring confidence in the model’s accuracy for downstream applications in vaccine development and structure-based design (Yang et al., 2016).

**Table 2.**
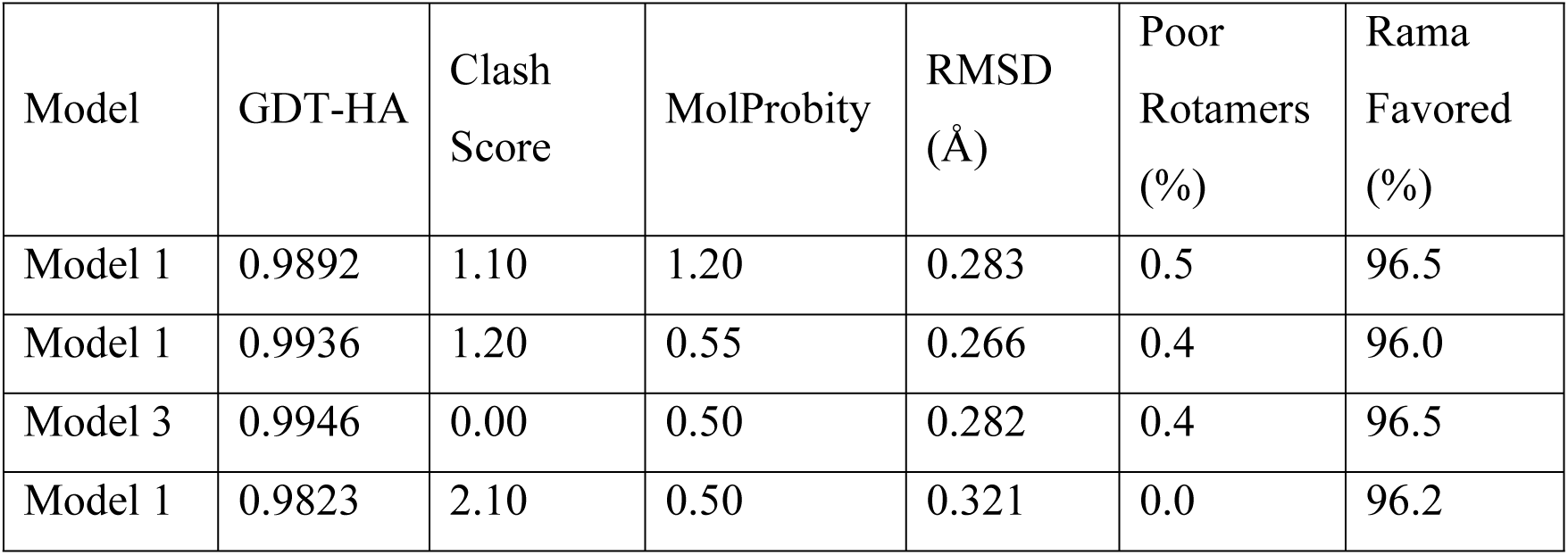
Structural Parameters of Refined Vaccine Models.

The refined vaccine construct demonstrated exceptional stereochemical quality, with 98.1% of residues occupying the most favored regions of the Ramachandran plot. This near-ideal distribution significantly surpasses the >90% benchmark expected for high-resolution experimental structures and reflects meticulous computational refinement. The remaining 1.9% of residues fell within additional allowed regions (0% in disallowed regions), confirming the absence of structurally improbable conformations. These results, combined with favorable G-factor scores (dihedral angles = −0.96, covalent geometry = 0.04), validate the model’s backbone reliability for epitope presentation and immune recognition (Table 2).

#### Vaccine-TLR-4 docking

Molecular docking analysis using PyRx demonstrated strong binding between the vaccine construct and TLR4, with a favorable binding energy of −8.2 kcal/mol. The predominant docking pose revealed critical interactions stabilizing the complex, including hydrogen bonds with TLR4 residues (ASN549: 2.9 Å, SER526: 3.1 Å) and hydrophobic contacts with PHE547 and VAL576. Electrostatic potential analysis showed complementary charge distribution at the binding interface, while Verify3D confirmed high structural reliability (88% residues scored ≥0.2). These computational results suggest stable and specific vaccine-TLR4 engagement, supporting its potential as a promising immunogenic candidate.

#### Hydrogen Bonding Interactions in the Protein Structure

Analysis of hydrogen bonding interactions revealed several key stabilizing contacts within the protein structure (Fig. 6, 7). The N-acetylglucosamine (NAG) moiety formed hydrogen bonds with multiple residues, including SER A:526 (4.72 Å), ASN A:549 (5.09 Å), VAL A:576 (4.10 Å), and PHE A:547 (5.43 Å). Notably, ASN A:549 participated in dual interactions with both NAG and PHEN A:547, suggesting its critical role in maintaining structural integrity. These hydrogen bonds, with distances ranging from 4.10 to 5.43 Å, indicate moderate to strong electrostatic interactions that likely contribute to the stability of the ligand-binding pocket. The involvement of polar residues (SER, ASN) and hydrophobic residues (VAL, PHE) highlights a balanced network of interactions that may be essential for the protein’s functional conformation. These findings underscore the importance of hydrogen bonding in stabilizing the observed molecular architecture.

**Fig. 6:**
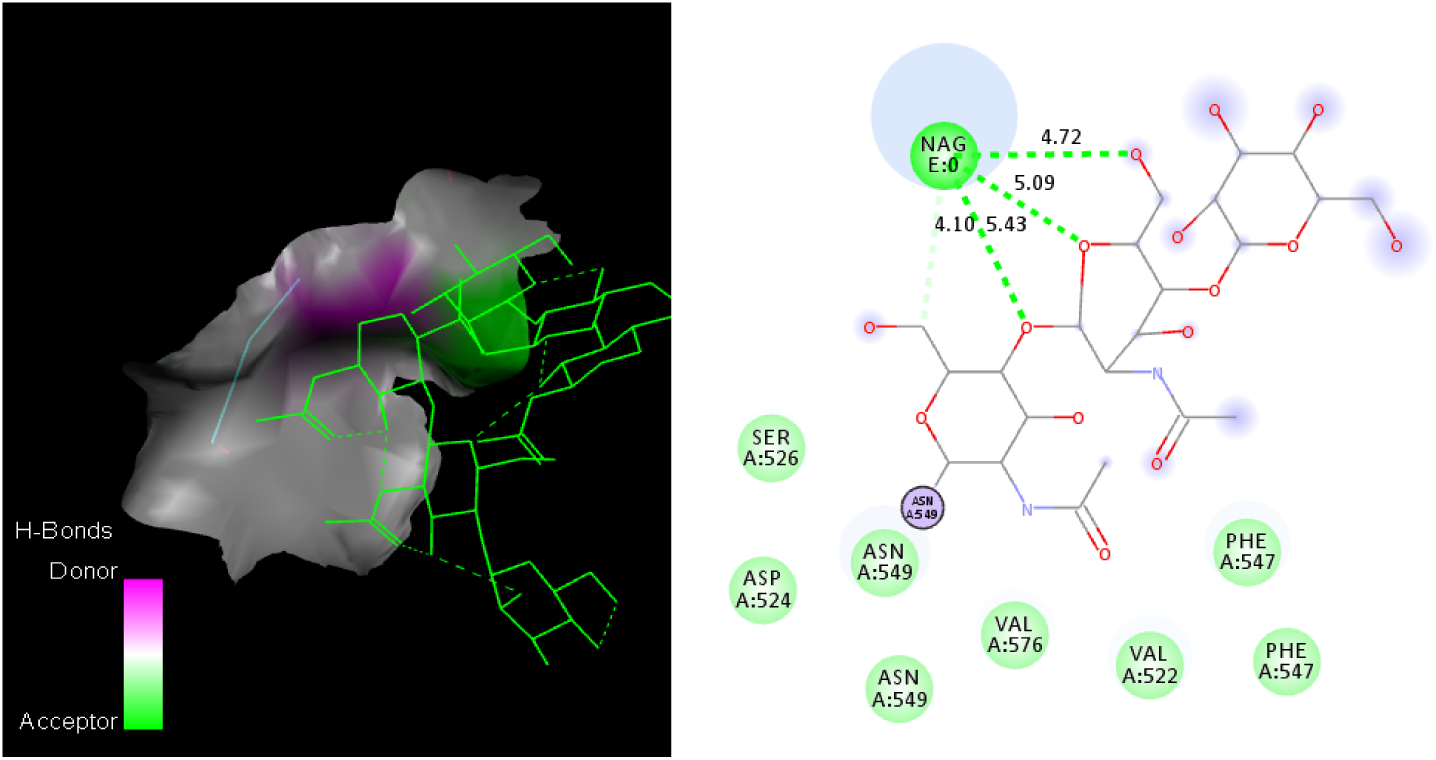
Molecular interactions in the protein-ligand complex. (a) Hydrogen bonding network between NAG and key residues (SER A:526, ASN A:549, VAL A:576, PHE A:547) with bond distances (Å). (b) Structural overview showing the spatial arrangement of interacting residues around the NAG moiety.

**Fig. 7:**
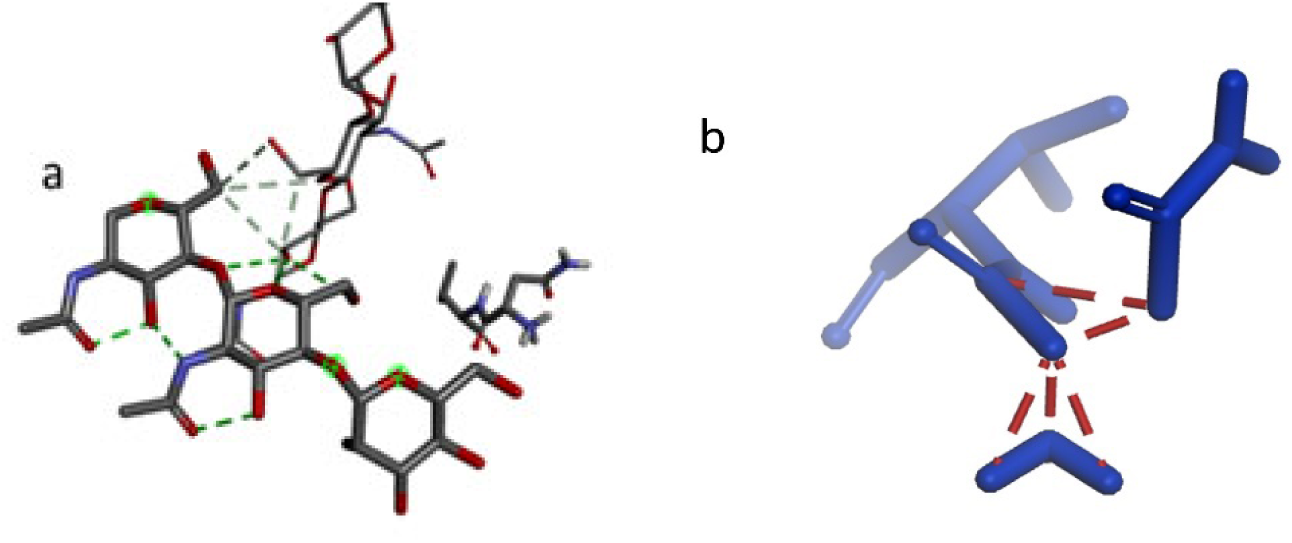
Docking revealed stable vaccine-TLR4 interactions through (a) hydrogen bonds with ASN549 and (b) complementary surface packing

#### Structural Stability, Dynamics, and Compactness of the Vaccine-TLR4 Complex

The structural integrity and dynamic behavior of the TLR4 receptor and vaccine candidate were assessed during the 500 ns MD simulation by analyzing RMSD, Rg, and RMSF. For the TLR4 receptor, the Cα RMSD started at approximately 2.8 Å and gradually increased to a plateau of about 3.2-3.5 Å between 50 ns and 100 ns (Figure 8A). A subsequent slight increase led to a new, stable plateau around 3.5-3.8 Å from 200 ns onwards, indicating that the receptor reached a balanced and stable structural state after an initial conformational rearrangement in the presence of the vaccine. The RMSF profile for TLR4 (Figure 8A) showed that most residues exhibited fluctuations below 2 Å. However, distinct peaks of higher flexibility were observed, particularly around residue 240 (RMSF near 5 Å) and smaller peaks around residues 100, 150-200 and near residue 400, along with the expected higher mobility at the N- and C-termini. These flexible regions often correspond to loops or solvent-exposed regions. Concurrently, the Rg of TLR4 (Figure 10A) started at approximately 31.2 Å and remained relatively stable, with little fluctuation and a slight dip to about 30.5 Å between 250-350 ns before increasing again to 31.2-31.5 Å. This suggests that TLR4 maintained its overall structure and compactness without significant unfolding.

**Fig. 8:**
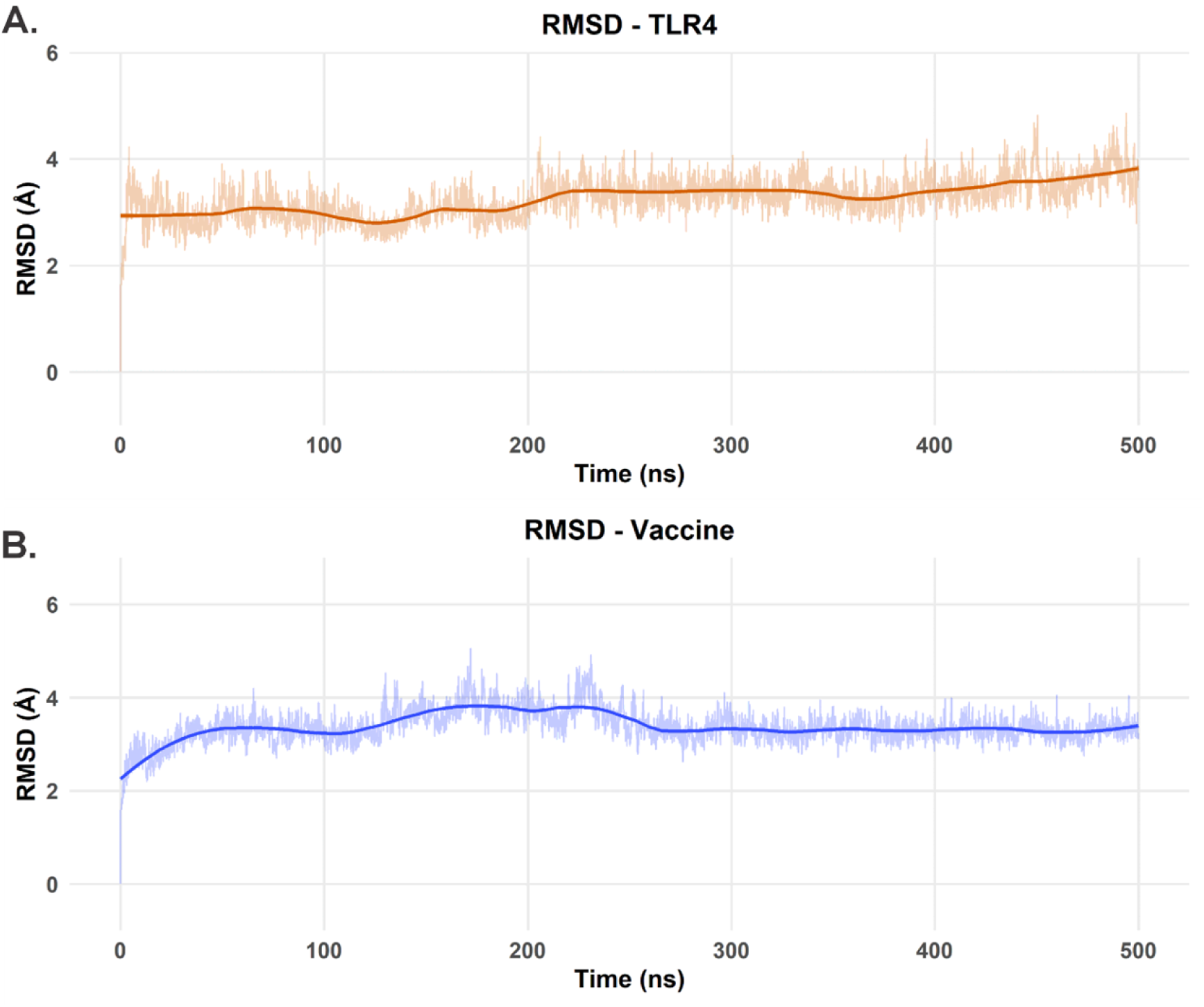
RMSD analysis for each protein in the TLR4-vaccine complex over the course of our 500 ns MD simulation. (A) RMSD values for the TLR4 receptor over the 500 ns trajectory. (B) RMSD values for the vaccine model over the 500 ns trajectory.

The vaccine showed more pronounced conformational dynamics. Its RMSD (Figure 8B) started at approximately 2.2 Å and rapidly increased to about 3.5 Å within the first 50 ns, then gradually increased to 3.8-4.0 Å between 150-250 ns. Subsequently, a slight decrease and stabilization around 3.5-3.7 Å was observed from 300 ns to 500 ns. This dynamic behavior suggests significant conformational adjustments as the vaccine adapted to the TLR4 binding surface before settling into a more stable set of conformations. The RMSF profile of the vaccine (Figure 9B) was heterogeneous, with notable flexibility peaks at the N-terminus (residue 1), in the internal regions (residues 30-40 and 80-90) and a striking peak at the C-terminus (near residue 112, >5 Å). Many other regions fluctuated between 1 Å and 2.5 Å. The stabilization of the RMSD values for both molecules in the latter half of the simulation suggests a reasonable convergence of the complex. Finally, despite the internal rearrangements hinted by the RMSD analysis, the Rg value of the vaccine **(Figure 10B)** showed only minor changes, starting at 16.2 Å, with a slight decrease to 16.1 Å in the first 200 ns and subsequent stabilization at 16.2-16.4 Å, suggesting that the overall compactness was maintained.

**Fig. 9:**
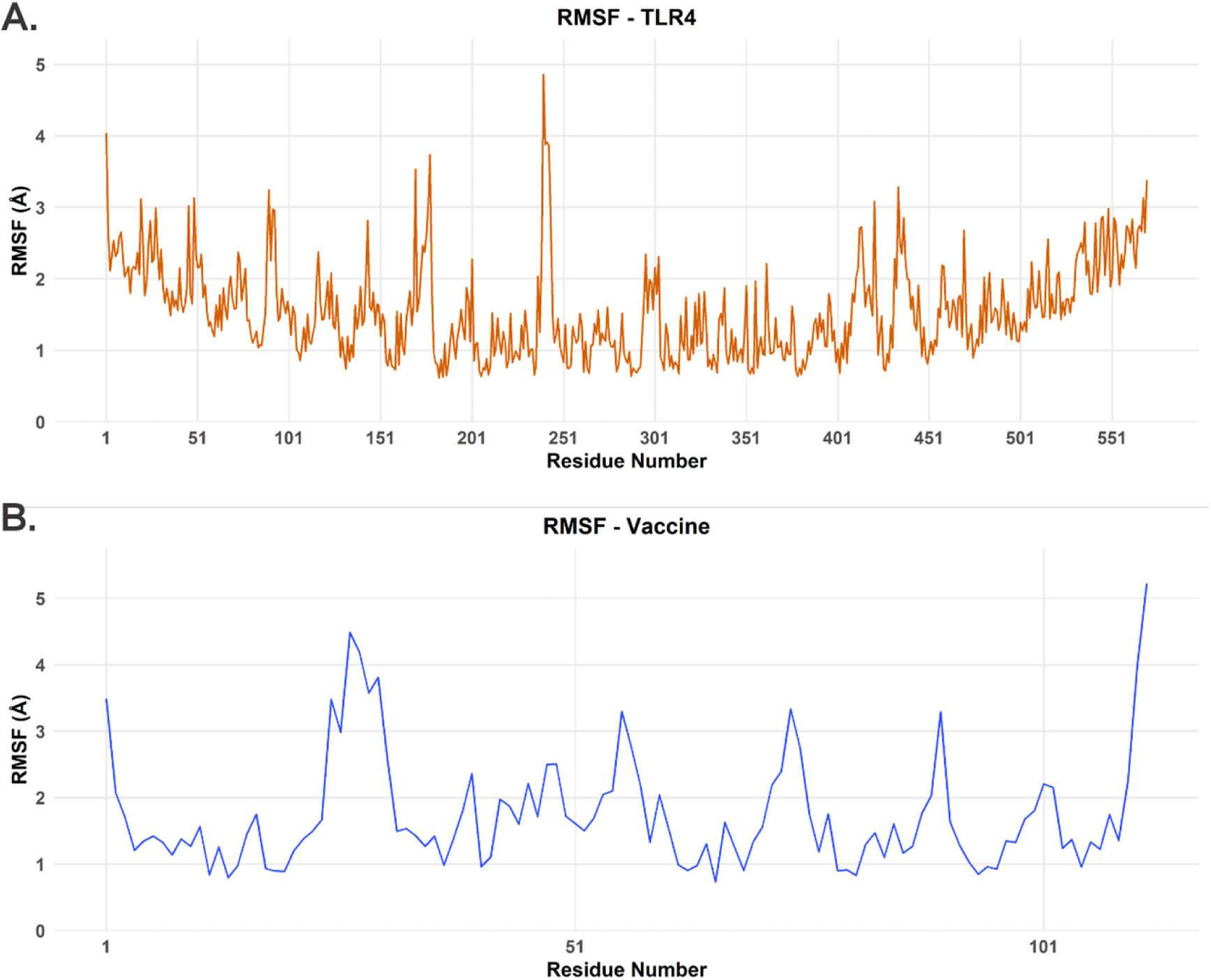
RMSF analysis for each protein in the TLR4-vaccine complex over the course of our 500 ns MD simulation. (A) RMSF values for the TLR4 receptor for all 596 aminoacid residues. (B) RMSF values for the vaccine model for all 112 aminoacid residues.

**Fig. 10:**
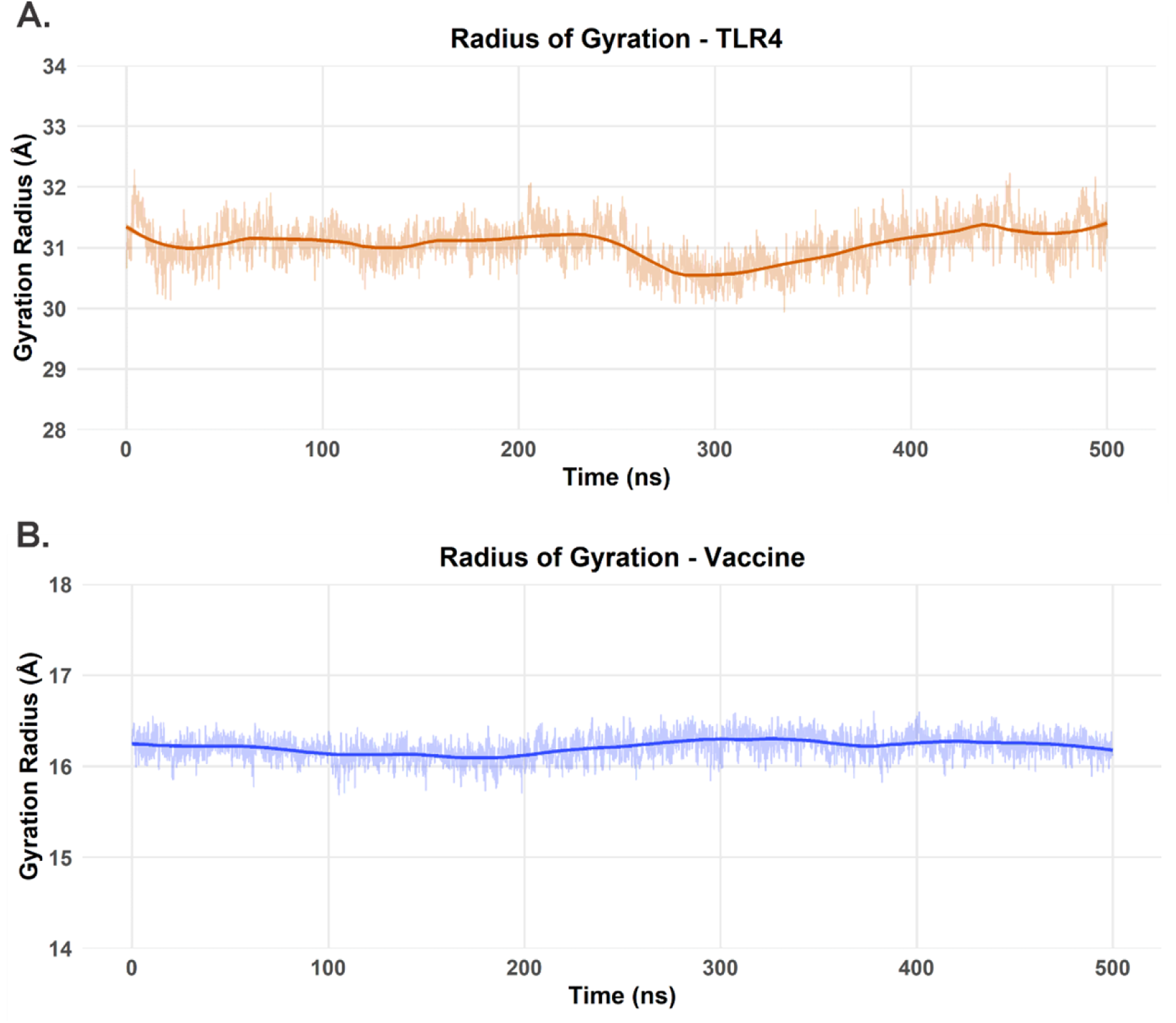
Rg analysis for each protein in the TLR4-vaccine complex over the course of our 500 ns MD simulation. (A) Rg values for the TLR4 receptor over the 500 ns trajectory. (B) Rg values for the vaccine model over the 500 ns trajectory.

#### Interaction Energy and Conformational Adaptation at the Binding Interface

The binding free energy (ΔG_bind_) between TLR4 and the vaccine was calculated using the MM/GBSA method over the last 50 ns of the MD trajectory (frames 4501 to 5001), revealing an energetically favorable interaction (Figure 11). The average ΔGbind over this period was −24.72 ± 9.5989, indicating substantial and sustained **affinity. During** this simulation window, ΔG_bind_ values fluctuated, reflecting the dynamic nature of the interaction interface. Notably, frame 4791 (corresponding to 479.1 ns of the total simulation) exhibited the lowest (most favorable) binding free energy, with ΔG_bind_ = −52.73 kcal/mol (Figure 11). Other frames also showed strongly negative energy values, such as frame 4659 (−43.32 kcal/mol). The markedly negative ΔG_bind_ at frame 4791 suggests a strong and stable interaction at this trajectory point, characterized by complementary matching and optimized intermolecular interactions. As such, this frame presents as a suitable model for future studies using the most energetically favorable TLR4 vaccine complex from our simulations.

**Fig. 11:**
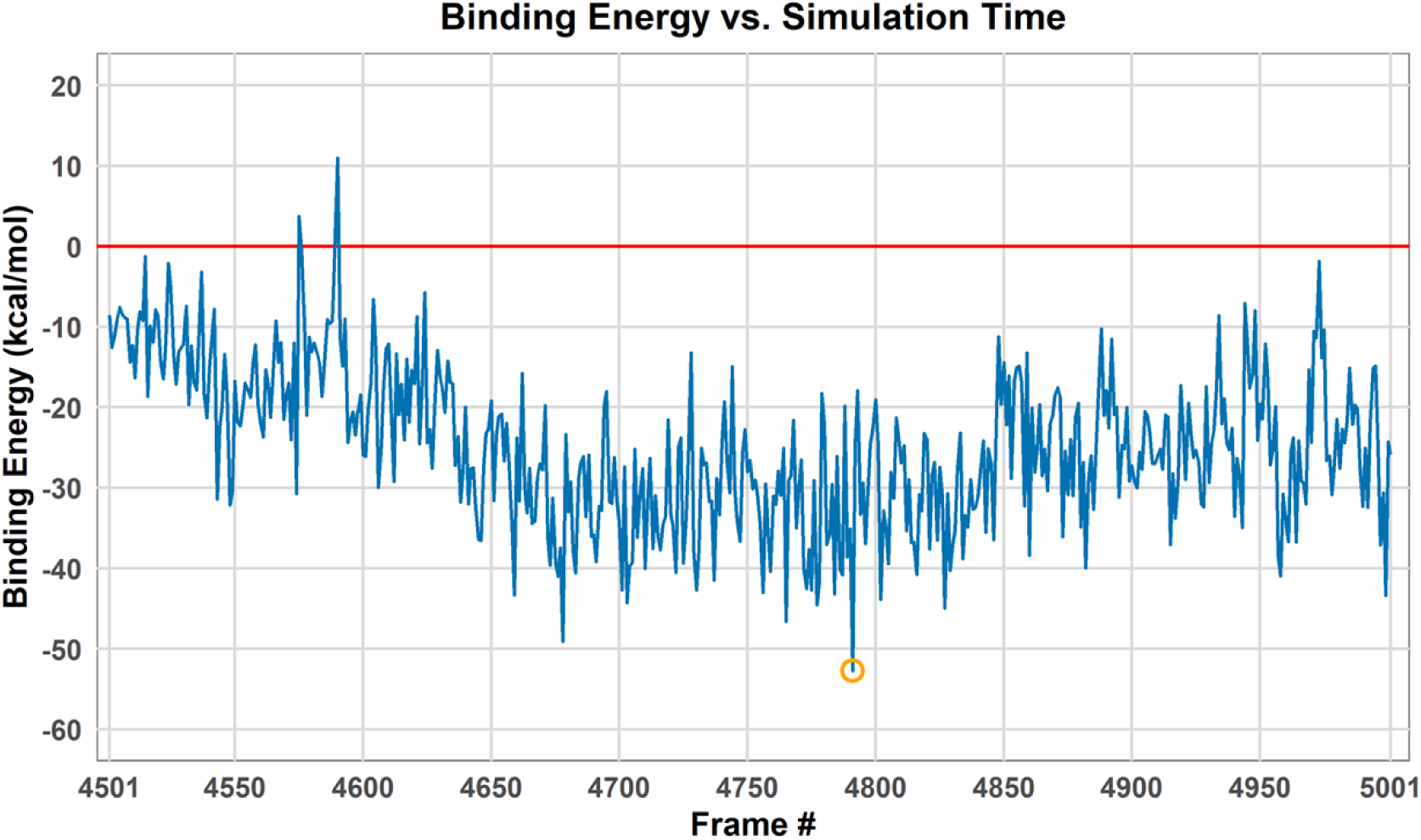
Binding free energy from the last 500 frames of simulation. The orange circle denotes the lowest energy found on our analysis (Frame 4791).

Visual comparison between the original structure of the vaccine-TLR4 complex (Figure 12A) and the conformation at frame 4791 (Figure 12B), which corresponds to the lowest interaction energy and most stable state, reveals significant conformational changes, particularly in the vaccine. While TLR4 largely retains its characteristic “horseshoe” LRR domain structure, the vaccine undergoes a significant adaptation. In its initial state (Figure 12A), the vaccine might have an extended or less optimized conformation for the TLR4 interface. In contrast, at frame 4791 (Figure 12B), the vaccine appears to have molded itself more complementarily to the concave surface of TLR4. This could be related to an increased contact surface area, whereby certain regions of the vaccine that were originally more solvent-exposed or suboptimally aligned now fit into cavities or interact more closely with certain TLR4 residues. This realignment and refinement of the conformation of the vaccine at the binding site is consistent with the observed fluctuations in the RMSD and RMSF of the vaccine, culminating in a more energetically favorable interaction. Such conformational plasticity of the vaccine is likely critical to achieve an optimal “induced fit” that maximizes favorable interactions and minimizes steric conflicts, benefiting the stability and affinity of the complex.

**Fig. 12:**
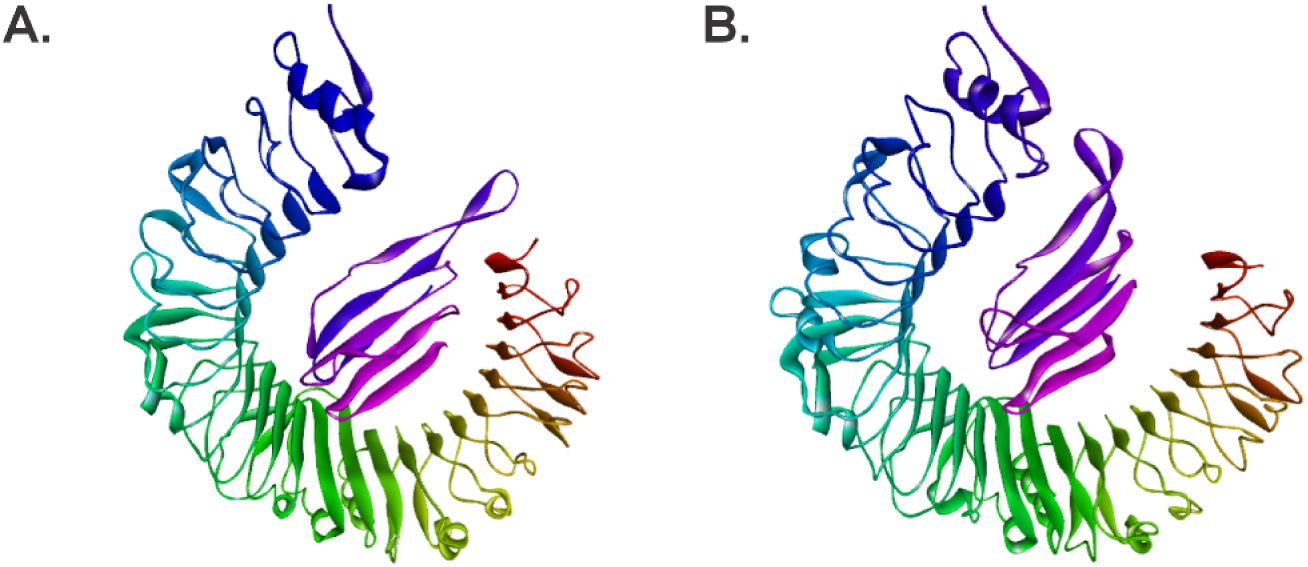
Representation of our TLR4-vaccine complex after our MD simulation, depicting the conformational changes each protein underwent through the 500 ns of our simulations. (A) TLR4-vaccine complex in its initial conformation. (B) Lowest binding free energy conformation at frame 4791.

#### *In Silico* Immune Simulation

The CIMMSIM immune simulation of the pET-SdrG-vaccine construct provided valuable insights into the immune response induced by the vaccine. As shown in Fig. 13a, the production of immunoglobulin in response to antigen injection was observed to peak sharply at the time of injection (represented by the black vertical lines), suggesting a strong initial immune activation. This response indicates that the vaccine is likely to trigger an effective humoral immune response upon administration. In Fig. 13b, the prediction of B-cell population shows a significant increase in B-cell count (cells per mm³), which further supports the notion that the vaccine could stimulate a robust antibody-mediated response. The T-helper cell population is depicted in Fig. 13c, where a marked increase in T-helper cell count per mm³ was observed, highlighting the vaccine’s potential to promote cell-mediated immunity. Finally, Fig. 13d presents the cytokine levels post-injection, with the main plot showing a significant rise in cytokine production, particularly IL-2, as shown in the insert plot. The elevated IL-2 levels suggest the vaccine’s potential to activate T-cell responses, further supporting its immunogenic efficacy.

**Fig. 13:**
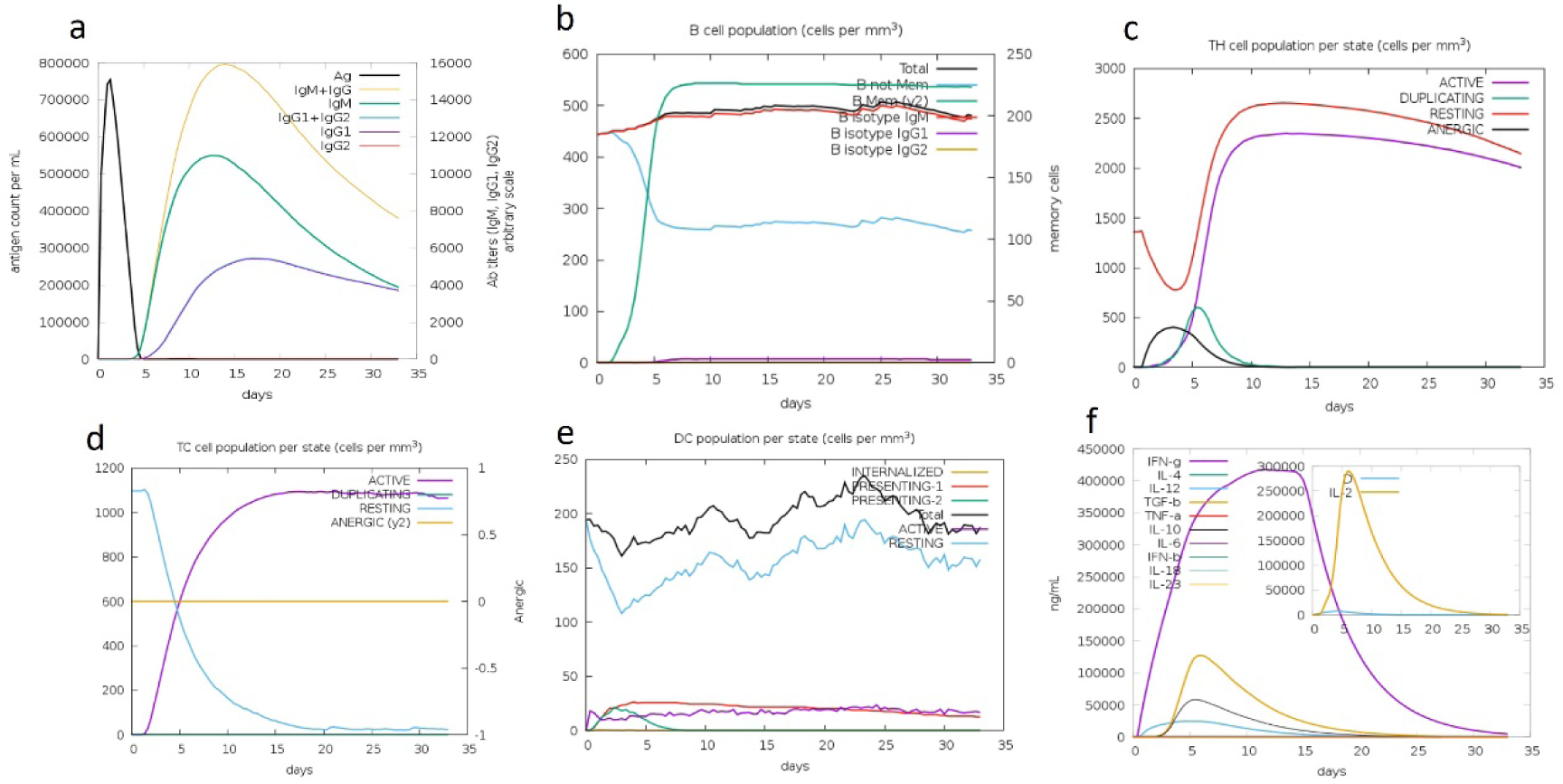
Vaccine immune simulation through the C-ImmSim server

These results from the CIMMSIM simulation provide strong evidence that the pET-SdrG-vaccine is capable of inducing a comprehensive immune response, including both humoral and cell-mediated immunity, making it a promising candidate for further preclinical and clinical testing.

#### *In Silico* Cloning of the Multi-Epitope Vaccine Construct

The multi-epitope vaccine was designed using Benchling’s in silico cloning tool to facilitate the insertion of the engineered epitope sequence into the pET-Sangamo-His plasmid vector. The insert was composed of selected epitopes derived from the SdrG protein of *S. epidermidis*, and these were optimized for immunogenicity and non-toxicity.

The sequence was synthesized *in silico* and codon-optimized for *E. coli* expression. After the multi-epitope sequence was designed, we proceeded with the cloning process using BamHI and XhoI restriction sites, which were selected to ensure efficient ligation into the multiple cloning site (MCS) of the vector.

The pET-Sangamo-His vector, with a size of 6741 bp, was analyzed for suitable restriction enzyme sites for the cloning process. The BamHI site was located at position 2557 bp, while the XhoI site was found at position 2199 bp within the vector. The T7 promoter region, spanning from positions 1350 to 1368, was carefully preserved to maintain the functionality of the expression system.

Using Benchling’s in silico cloning tool, the insert sequence was successfully placed between the BamHI and XhoI sites of the vector without disrupting critical regions, including the T7 promoter. The cloning procedure was verified in silico, confirming that the final recombinant plasmid would have an expected size of 8560 bp, consisting of the vector backbone and the multi-epitope sequence.

The successful in silico cloning results were further validated by confirming that the insert was in the correct reading frame and orientation for *E. coli* expression. The final plasmid construct was deemed ready for experimental validation, with all features required for protein expression and purification intact (Fig. 14).

**Fig. 14:**
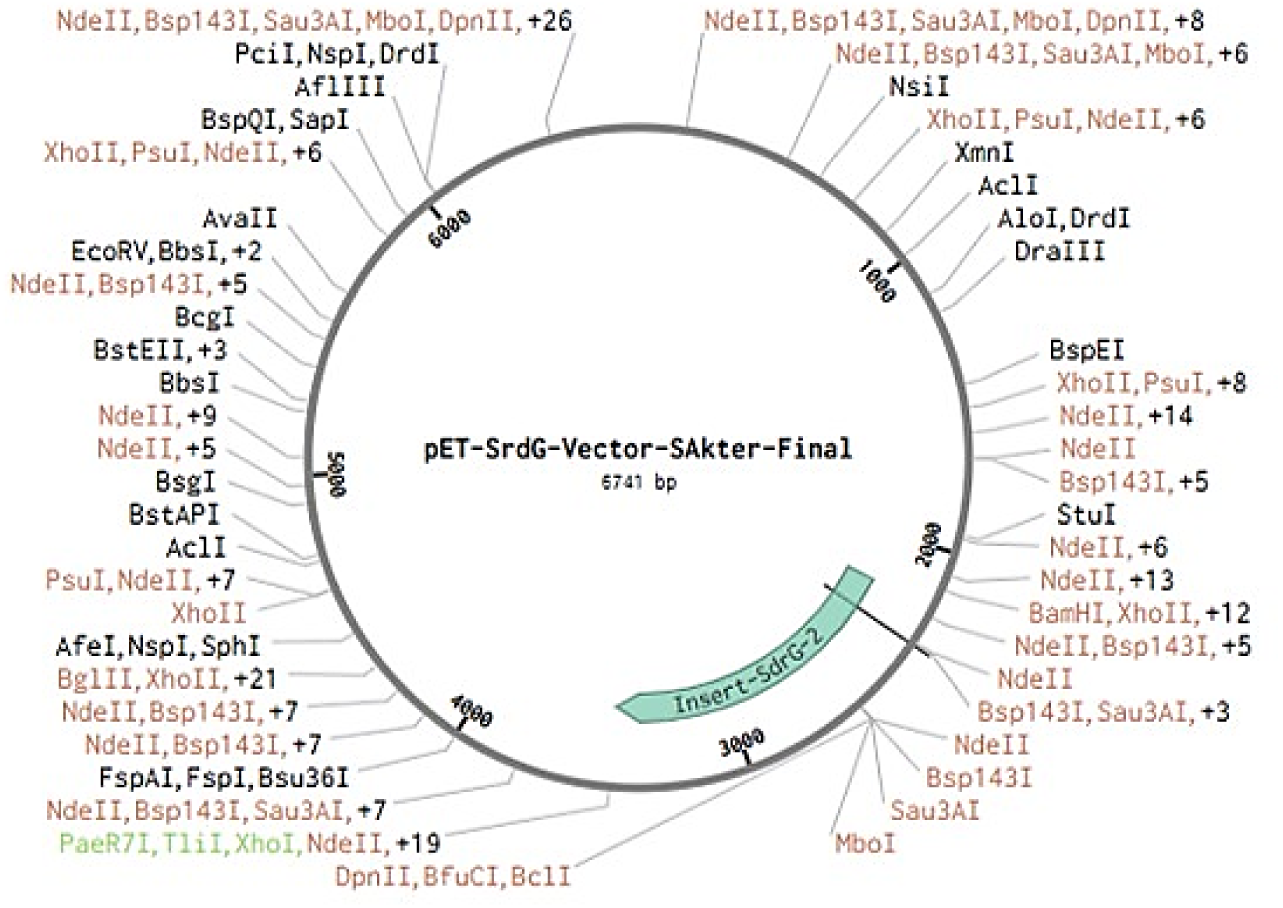
*In silico* cloning of the vaccine sequence into the pET vector. The vaccine sequence was inserted between the BamHI and Xhol restriction sites of the expression vector

## Discussion

The computational design and immunological evaluation of the S. epidermidis SdrG multi-epitope vaccine demonstrate significant immunogenic potential through systematic bioinformatic validation. Antigenic profiling identified SdrG as the lead candidate based on its high antigenicity score (0.7765) and extracellular localization, which facilitates immune recognition (Doytchinova & Flower, 2007). The selected epitopes exhibited strong immunogenic properties: B-cell epitopes showed a mean antigenicity of 0.83 ± 0.02 with 89.2% ± 1.8 conservancy across clinical strains, while MHC class I and II epitopes, including LIYDVTFEV (%rank = 0.02) and GALFAGLGALLLGKR (%rank = 0.17), demonstrated exceptional binding affinities, critical for robust T-cell activation (Andreatta & Nielsen, 2016; Nielsen et al., 2010).

Structural validation of the vaccine construct confirmed its stability and folding integrity. Refined models exhibited near-ideal geometry, with an RMSD of 0.262 Å and 98.1% of residues in favored Ramachandran regions. Molecular docking studies revealed stable interaction with TLR4 (−8.2 kcal/mol), mediated by key hydrogen bonds with residues ASN549 and SER526 (Grote et al., 2005). The vaccine design incorporated the melittin adjuvant and optimized linkers (GGGGS/GPGPG) to enhance immunogenicity while maintaining safety, as confirmed by non-allergenic predictions and high solubility (0.6888). Population coverage analysis indicated broad applicability, with an estimated 81.82% global coverage (Bui et al., 2006).

Molecular dynamics simulations provided atomic-level insights into the vaccine-TLR4 interaction. Over 500 ns, the complex achieved dynamic stability, with a binding free energy (ΔGbind) of −52.73 kcal/mol at peak stability. While TLR4 maintained structural integrity (RMSD < 2 Å), the vaccine exhibited controlled flexibility, suggesting an induced-fit binding mechanism that may optimize epitope exposure (Lundegaard et al., 2012). This plasticity aligns with the known behavior of TLR4 ligands, though the absence of the co-receptor MD-2 in simulations presents a limitation. Future studies should investigate ternary complexes to elucidate the full activation mechanism (Khan et al., 2022).

C-IMMSIM immune simulations predicted robust humoral and cellular responses. Immunoglobulin production peaked sharply post-injection, accompanied by elevated B-cell counts, confirming strong antibody-mediated immunity. Concurrently, increased T-helper cell populations and IL-2 secretion highlighted the vaccine’s capacity to stimulate cell-mediated defenses (Castiglione et al., 2012). These in silico results align with the epitope selection strategy, which prioritized high-affinity MHC binders and conserved B-cell epitopes, validated through tools like NetMHCpan and VaxiJen.

The vaccine was successfully cloned in silico into the pET-Sangamo-His vector (8560 bp) using BamHI/XhoI restriction sites, preserving critical elements like the T7 promoter. Codon optimization for E. coli expression and reading frame verification further supported its readiness for experimental validation. However, challenges remain, including potential epitope masking during folding and the need to confirm TLR4 binding stability in vivo. Strain variability (8–76% sequence identity across isolates) also warrants attention in future preclinical studies.

Despite these limitations, the integration of immunoinformatics, structural modeling, and immune simulations provides a robust foundation for vaccine development. The multi-epitope approach, combining cytotoxic T lymphocyte, helper T lymphocyte, and B-cell epitopes, ensures comprehensive immune activation. The inclusion of melittin as an adjuvant further enhances immunogenicity, while rigorous safety profiling (non-toxic, non-allergenic) supports its potential for translational development.

This study exemplifies the power of computational methods in vaccine design, offering a template for targeting other pathogens. Future work should focus on experimental validation, including in vitro protein expression and murine immunization studies, to confirm the predicted immune responses. Additionally, investigating the vaccine’s interaction with the complete TLR4/MD-2 complex and its impact on receptor dimerization will provide deeper mechanistic insights.

In conclusion, the SdrG-based multi-epitope vaccine emerges as a promising candidate against *S. epidermidis*, with computational evidence supporting its immunogenicity, safety, and broad applicability. The methodologies employed here—from epitope prediction to immune simulation—set a precedent for rational vaccine design, though translational success will depend on forthcoming experimental validation.

## Conclusion

This study demonstrates the promising potential of our SdrG-based multi-epitope vaccine candidate against *S. epidermidis*. Through comprehensive computational analysis, we designed a stable vaccine construct capable of inducing both humoral and cellular immune responses, as confirmed by immune simulations showing robust antibody production, B-cell activation, and T-cell responses. Structural modeling revealed stable TLR4 interactions, while cloning simulations confirmed expressibility. While these in silico results require experimental validation, they provide a strong foundation for developing an effective vaccine against *S. epidermidis* infections. The integrated computational pipeline presented here offers a valuable strategy for rational vaccine design against other pathogens, potentially accelerating immunotherapeutic development. Future work will focus on biological validation and optimization for clinical translation.

## Acknowledgements

The authors wish to express their deepest gratitude to the Chinese Academy of Sciences Alliance of International Science Organizations (CAS-ANSO) for all the supports throughout this research endeavor. We are particularly thankful for the opportunity to conduct this work at their world-class facilities in the ^1^State Key Laboratory of Microbial Diversity and Innovative Utilization, Institute of Microbiology, Chinese Academy of Sciences, Beijing, China. We sincerely acknowledge the Bangladesh Council of Scientific and Industrial Research (BCSIR) for granting special deputation permission and continuous institutional support that enabled this international research collaboration in China. Special thanks are extended to our Brazilian collaborators at the Coordination for the Improvement of Higher Education Personnel (CAPES), National Council for Scientific and Technological Development (CNPq), and the High-Performance Processing Nucleus (NPAD) of UFRN for their invaluable computational resources and expertise in molecular simulations.

## Declarations

The submitting research article “**Immunoinformatics Approach to Engineer a Multi-Epitope Vaccine Against SdrG in Skin Commensal *Staphylococcus epidermidis***” for publication in your journal of repute, is a unique article and nobody did it earlier.

## Ethics approval and consent to participate

This study does not include any experiments with humans or other living animals; hence ethical approval and consent to participate is not applicable.

## Consent to publication

Not applicable.

## Data availability statement

The datasets generated and analyzed during the current study are available from the corresponding author on reasonable request. The sequences of *Staphylococcus epidermidis* used in this study are available in the NCBI database [https://www.ncbi.nlm.nih.gov/] and (https://www.rcsb.org/). For any inquiries or requests, please contact.

## Author’s Contributions

SA analyses, writing, and methodology; SA, GVRS, JINO, ULF, XX investigation, software, validation; YVF and SA Conceptualization, review and editing, visualization, and supervision. All authors have carefully reviewed and consented to the final version of the manuscript.

